# Synergistic Multi-Enzyme Cascades Assembled on Modular Protein Scaffolds for Efficient PET Biorecycling

**DOI:** 10.1101/2024.09.16.613172

**Authors:** Yujia Zhang, Chongsen Li, Ehsan Hashimi, Enting Xu, Xuemei Yang, Yanbing Lin, Hui Gao, Zhuobin Liang

## Abstract

The global crisis of polyethylene terephthalate (PET) waste demands new strategies for sustainable management. Biocatalytic recycling offers a promising route, but direct depolymerization under moderate conditions is often limited by kinetic constrains of individual enzyme. Here, we engineer and optimize modular protein scaffolds that utilize orthogonal interaction domains, including the cohesin-dockerin pairs from bacterial cellulosomes and the SH3 ligand/domain pair from mouse Crk, to assemble complementary hydrolases (PETase, MHETase, and ICCG) into a synergistic multi-enzyme cascade. This architecture overcomes intermediate inhibition, efficiently converting PET hydrolysis products into the final monomers, terephthalic acid (TPA) and ethylene glycol (EG), outperforming previous multi-enzyme systems. The system’s application is further evaluated by stability enhancement via immobilization on metal-organic frameworks (MOFs), coupling depolymerization to the valorization of EG into glycolic acid (GA), and adapting the system for scalable whole-cell biocatalysis. This scaffold-based multi-enzyme approach provides a new strategy for developing integrated biocatalytic systems to advance a circular PET economy.

## Main

Polyethylene terephthalate (PET), a ubiquitous polyester valued for its durability and versatility, poses a major sustainability crisis due to its environmental persistence and the limitations of traditional recycling. Its vast global production, exceeding 30 million tons annually, leads to widespread plastic pollution, harming terrestrial and marine ecosystems ^1^, releasing hazardous microplastics ^2^, and posing risks to human health ^3^. Traditional mechanical recycling requires extensive sorting, often yields low-value products due to incomplete degradation ^4^, and can further exacerbate microplastic pollution ^5^. Chemical recycling routes such as methanolysis and glycolysis, while promising, typically require high-temperature operations and the complex purification steps to isolate the final products ^6,7^.

Biocatalytic degradation, harnessing enzymes from microorganisms to break down PET polymers, has been proposed as an alternative strategy ^8^. This approach is attractive due to its potential for complete degradation, lower environmental impact, and the generation of valuable products through recycling and upcycling, as showcased in early-phase industrial demonstrations ^9^. Several PET hydrolases, including TfH ^10^, leaf-branch cutinase (LCC) ^11^, and *Is*PETase ^12^, have been identified and further engineered ^13,14^ (**Supplementary Table 1**). However, existing biocatalytic systems still face major hurdles such as incomplete monomer production ^15^, intermediate inhibition ^16^, limited efficiency on high-crystallinity PET, and cost-effectiveness ^17^, hindering their widespread industrial implementation.

Recent efforts to address these limitations have focused on multi-enzyme systems that leverage the complementary action of different PET hydrolases ^18–20^ (**Supplementary Table 2**). While free enzyme mixtures successfully alleviate intermediate accumulation, their synergistic efficiency is constrained by stochastic diffusion and a lack of precise spatial organization ^19–20^. Conversely, while direct genetic fusions successfully enforce spatial proximity ^18^, these static single-chain architectures are structurally challenging to expand or modify. Consequently, current approaches generally lack the structural modularity required to easily substitute enzyme variants, integrate additional functional modules, and readily adapt the cascade for diverse applications like monomer upcycling and whole-cell catalysis.

To address these challenges, here we engineer a modular, self-assembling multi-enzyme system, inspired by the natural cellulosome architect. This system enables the precise co-localization of multiple hydrolases, creating a synergistic complex that overcomes the kinetic bottlenecks of individual enzymes. We term this system the Scaffold-enabled PET Enzyme Ensemble-augmented Degradation platform (SPEED). We report the iterative design and optimization of the system, followed by demonstrations of its industrial relevance through enhanced stability via immobilization, direct coupling of depolymerization to monomer valorization, and adaptation for whole-cell biocatalysis. This work establishes a scaffold-based framework for engineering efficient, multi-step biocatalytic pathways for PET plastic recycling and upcycling.

## Results

### Synergistic enzyme-scaffold system for enhanced PET degradation

To harness the complementary properties of different PET hydrolases, we designed a multi-enzyme system based on the two-step PET hydrolysis pathway (**Fig. 1a**). Building on previous work that demonstrated the synergy of combining PETase and MHETase ^18^, we selected FAST-PETase ^14^ (FAST) for its high catalytic activity and MHETase ^12^ for its ability to convert inhibitory intermediates bis-(2-hydroxyethyl) terephthalate (BHET) and mono-(2-hydroxyethyl) terephthalate (MHET) into the final monomers, terephthalic acid (TPA) and ethylene glycol (EG).

**Fig. 1:**
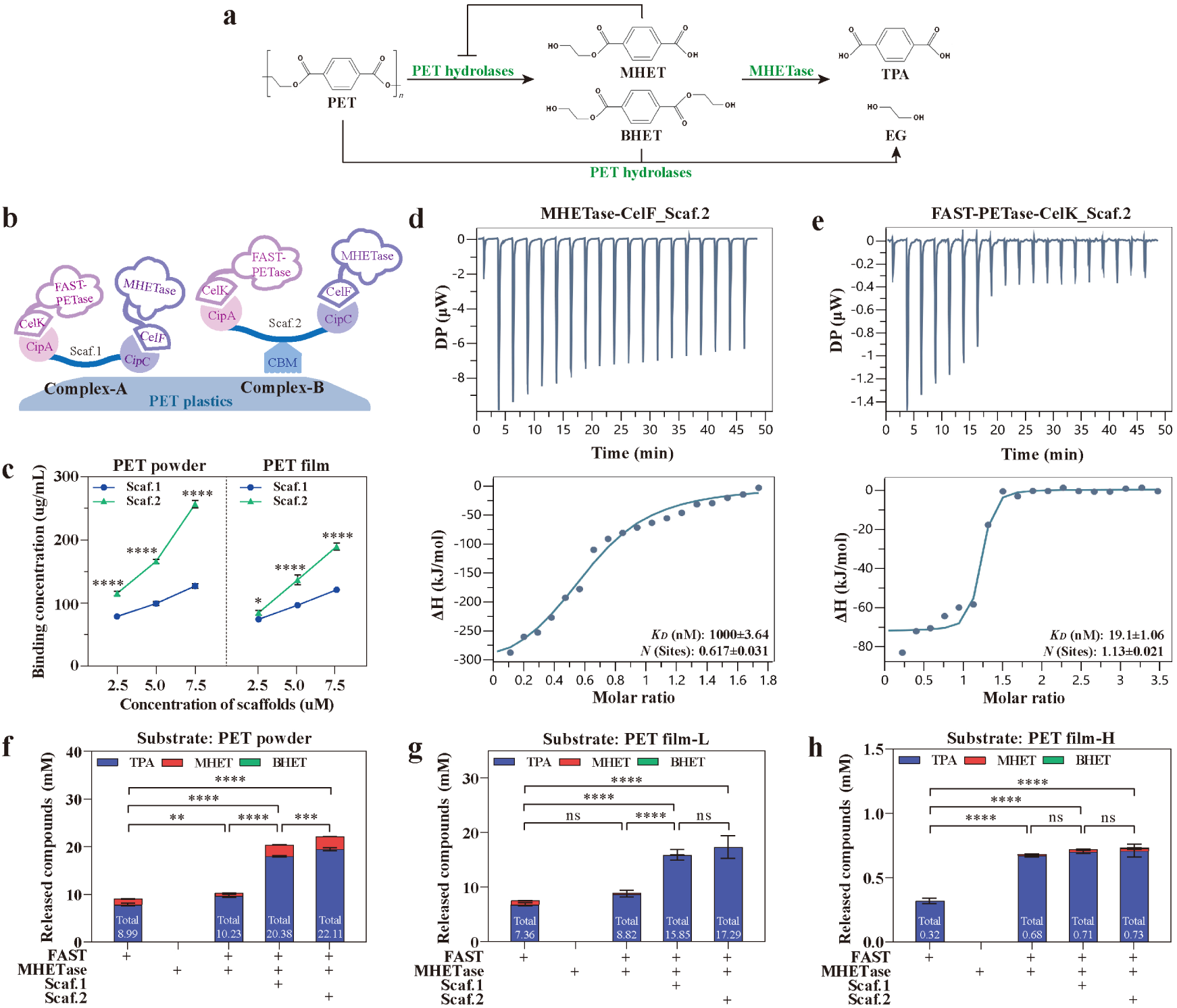
Scaffold assembly of PET hydrolases enhances depolymerization. **a** Schematic of the two-step PET hydrolysis pathway. PET is hydrolyzed by PET hydrolases to intermediates (MHET, BHET), which are subsequently converted to monomers (TPA, EG) by MHETase. **b** Schematic of the initial scaffold design. Dockerin-fused enzymes self-assemble onto cohesin-containing scaffolds to form Complex-A (Scaf.1) and Complex-B (Scaf.2). CelF/CipC: orthogonal dockerin/cohesin from *Clostridium cellulolyticum*; CelK/CipA: orthogonal dockerin/cohesin from *Clostridium thermocellum*. **c** Binding assay showing the affinity of Scaf.1 and Scaf.2 for PET powder and film substrates. **d**, **e** Isothermal titration calorimetry (ITC) confirms the binding of MHETase-CelF (**d**, K_D_ = 1000 nM) and FAST-PETase-CelK (**e**, K_D_ = 19.1 nM) to Scaf.2. **f–h** PET degradation assays comparing free enzymes (FAST, MHETase) with scaffolded complexes (Complex-A, Complex-B) on PET powder (**f**), low-crystallinity film (film-L) (**g**), and high-crystallinity film (film-H) (**h**). Reactions contained 100 mg PET, 0.1 nmol of each enzyme with an enzyme:scaffold molar ratio of 0.2:1, were incubated at 30 °C, pH 7.0, for 96 h. Data are presented as mean ± s.d. (n = 3). Statistical significance was determined by one-way ANOVA (ns, not significant; *P* < 0.05, \**P* < 0.01, \*\**P* < 0.001, \*\*\**P* < 0.0001).

Inspired by natural cellulosomes ^21,22^, we engineered and purified dockerin-fused versions of these enzymes to self-assemble onto cohesin-containing scaffolds, Scaf.1 and Scaf.2 (**Supplementary Fig. 1** and **2**), forming Complex-A and Complex-B, respectively (**Fig. 1b**). Scaf.2 includes a carbohydrate-binding module (CBM) ^23^ to enhance affinity for the PET substrates, which was confirmed in a binding assay (**Fig. 1c**). Isothermal titration calorimetry (ITC) further verified the binding of both MHETase-CelF and FAST-PETase-CelK to the Scaf.2 scaffold (**Fig. 1d,e**).

PET biodegradation assays confirmed the functional synergy of the assembled complexes in hydrolyzing PET substrates of varying crystallinity. These substrates, including PET powder, a low-crystallinity PET film (film-L), and a high-crystallinity PET film (film-H), sourced from suppliers common to the field and their crystallinity were further characterized by differential scanning calorimetry to ensure comparability with previous work (**Supplementary Fig. S3**). While combining free FAST and MHETase enhanced degradation compared to FAST alone, assembling them onto the scaffolds further boosted performance (**Fig. 1f-h**). Complex-B (using Scaf.2 with a CBM module) slightly outperformed Complex-A, yielding 22.11 mM (PET powder), 17.29 mM (film-L), and 0.73 mM (film-H) total products after 96 hours.

While these results are consistent with the synergy benefit reported in an earlier dual-enzyme fusion system ^18^, our modular scaffold provides the structural flexibility required to expand the multi-enzyme cascade and achieve improved degradation performance (**Supplementary Table 2** and **File 1-2**).

### Optimization of MHETase expression, assembly, and activity

Despite the above validated benefits of MHETase, the relatively low expression of recombinant MHETase in *E. coli* hindered further development. To address this bottleneck, we pursued a multi-pronged optimization strategy focusing on host strain selection, induction expression conditions, codon usage, and protein domain fusions, that improved MHETase expression and purification in *E. coli* (**Fig. 2a**, **Supplementary Fig. 4** and **5**). We replaced its fused CelF dockerin with a Src Homology 3 (SH3) peptide ligand (SH3_L_) and redesigned the scaffold (Scaf.3) to include the corresponding SH3 domain (SH3_D_) from the mouse Crk protein ^24^ for assembly (**Fig. 2b**). We discovered that the fusion topology was critical for this new interaction: an N-terminal fusion of the SH3_L_ to optiMHETase resulted in high-affinity binding to Scaf.3 (K_D_ = 12.6 nM), whereas a C-terminal fusion abrogated binding entirely (**Fig. 2c,d**). This optimized N-terminal construct achieved an ∼ 80-fold enhancement in binding affinity over the original CelF dockerin-based system (**Fig. 2c** vs. **Fig. 1d**), and was comparable to the FAST-PETase-CelK interaction with Scaf.3 (**Fig. 2e**). These successful modifications enabled the assembly of the improved Complex-C (**Fig. 2b**).

**Fig. 2:**
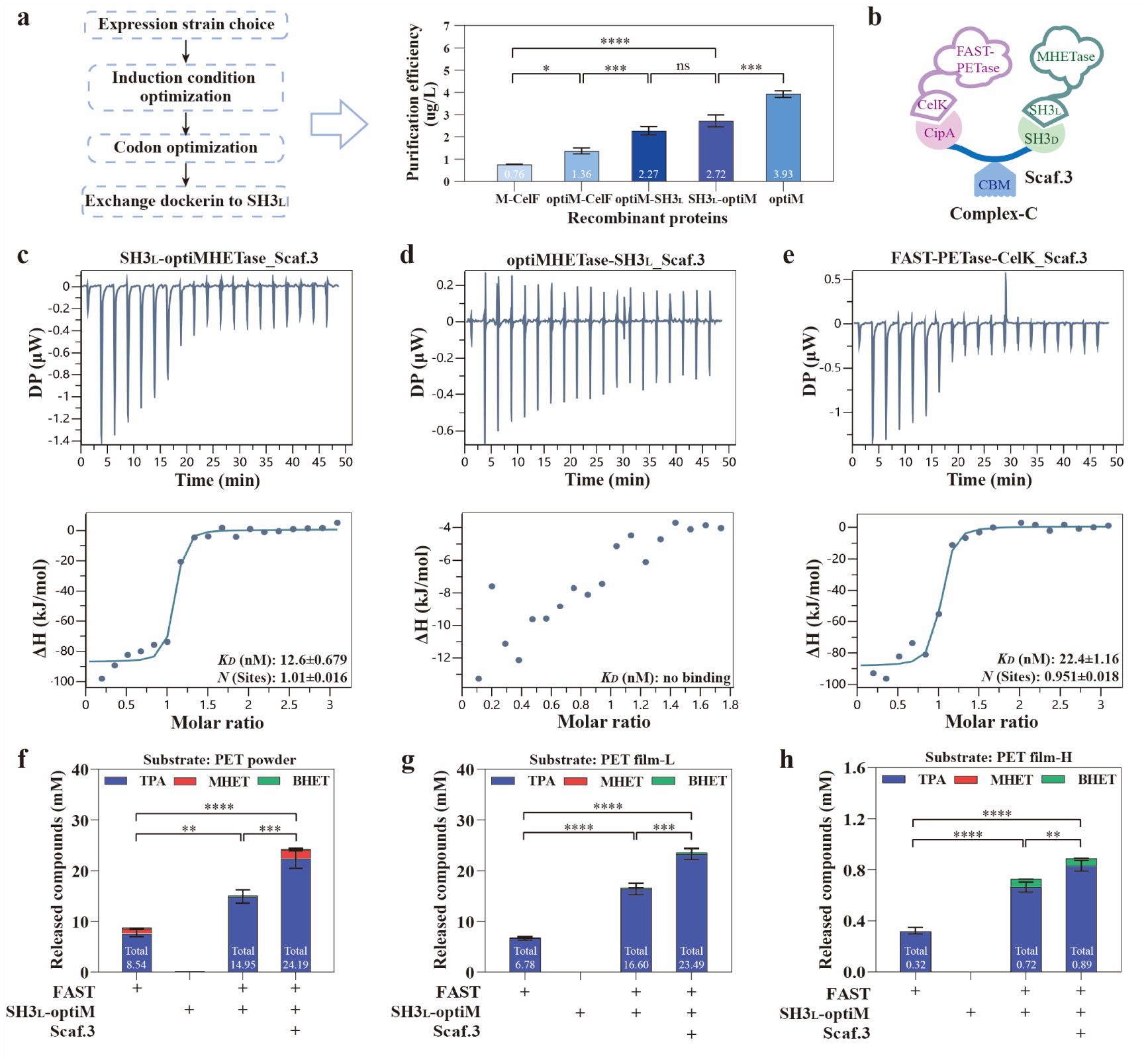
Optimization of MHETase enhances scaffold assembly and complex activity. **a** Schematic illustration of MHETase optimization strategy and the resulting purification efficiency of different MHETase variants from selected *E. coli* C41(DE3) strain with optimal induction condition (16 °C with 1 mM IPTG for 18 h). Purification efficiency is defined as the final yield of purified protein (ug) from each liter (L) of *E. coli* cultures with similar OD (cell density). M-CelF: MHETase-CelF; optiM-CelF: codon optimized M-CelF; M-optiSH3_L_: optiMHETase-SH3_L_; SH3_L_-optiM: SH3_L_-optiMHETase; optiM: codon optimized MHETase (as control). **b** Schematic of the redesigned Complex-C, assembled on the Scaf.3 (CipA-CBM-SH3_D_). SH3_D_/SH3_L_: SH3 domain sourced from the mouse protein Crk and its cognate peptide ligand. **c**-**e** ITC binding analysis of SH3_L_-optiMHETase **(c)**, optiMHETase-SH3_L_ **(d)**, and FAST-PETase-CelK **(e)** to the redesigned scaffold Scaf.3. **f**-**h** PET degradation assays comparing individual enzymes, enzyme mixtures, and the fully assembled Complex-C (FAST + SH3_L_-optiM + Scaf.3), on PET powder **(f)**, film-L **(g)**, and film-H **(h)**. SH3_L_, SH3 ligand; SH3_D_, SH3 domain. Reactions contained 100 mg PET, 0.1 nmol of each enzyme with an enzyme:scaffold molar ratio of 0.2:1, were incubated at 30 °C, pH 7.0, for 96 h. Data are presented as mean ± s.d. (n = 3). Statistical significance was determined by one-way ANOVA (ns: not significant, *p < 0.1, **p < 0.01, ***p < 0.001, ****p < 0.0001).

Degradation assays across various PET substrates revealed significant improvements (**Fig. 2f-h**). Complex-C yielded 22.37 mM, 23.22 mM and 0.83 mM TPA production with PET powder, film-L and film-H, respectively. This represents a 3-fold increase over FAST-only groups and a 22.2% average increase over Complex-B (**Fig. 2f-h** vs. **Fig. 1f-h**). These results confirm Complex-C’s effectiveness in promoting PET recycling.

### Modular scaffold expansion to boost PET degradation capacity

The transition from Complex-A to Complex-C demonstrates the modularity of the SPEED design, allowing for tailored enzyme combinations for diverse substrates. To broaden substrate range, we next incorporated ICCG, an enzyme active towards higher-crystallinity substrates (6%-40%) ^25^, complementing the preference of FAST-PETase for lower-crystallinity PET plastics (1%-30%) ^14^. Using the dockerin-cohesin pair ScaA-ScaB ^26^, ICCG was integrated into a new scaffold, CipA-CBM-SH3_D_-ScaB (Scaf.4), forming Complex-D (**Fig. 3a** and **Supplementary Fig. 6**).

**Fig. 3:**
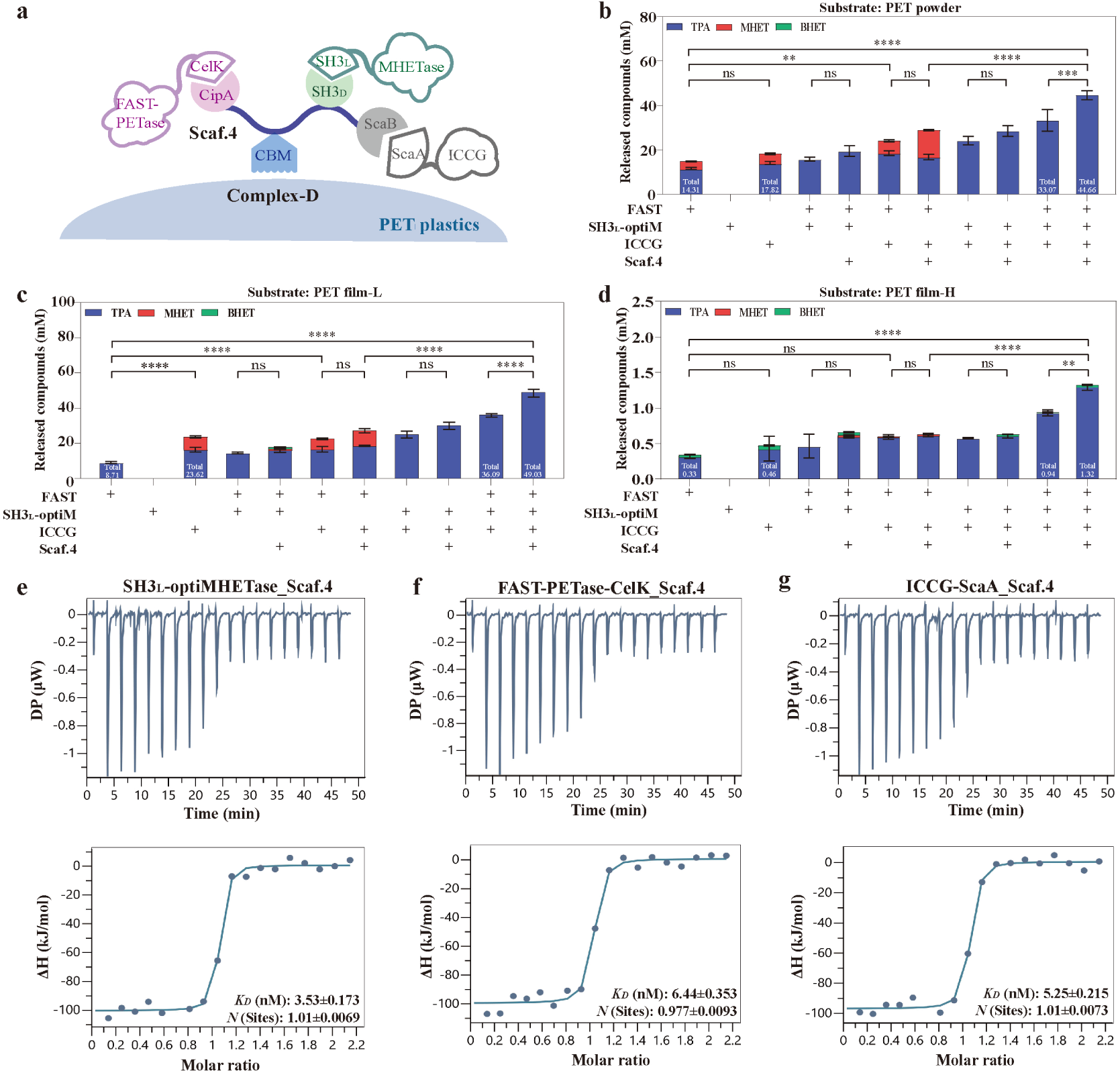
A three-enzyme scaffolded cascade improves PET degradation and monomer purity. **a** Schematic of the expanded three-enzyme complex (Complex-D), incorporating FAST-PETase, optiMHETase, and ICCG on the modular Scaf.4 scaffold. ScaA/ScaB: orthogonal dockerin/cohesin pair from *Ruminococcus flavefaciens*. **b**-**d** PET degradation assays on PET powder (**b**), film-L (**c**), and film-H (**d**) comparing individual enzymes, enzyme mixtures, and the fully assembled Complex-D (FAST + SH3L-optiM + ICCG + Scaf.4). Reactions contained 100 mg PET, 0.1 nmol of each enzyme with an enzyme:scaffold molar ratio of 0.2:1, were incubated at 30 °C, pH 7.0, for 96 h. **e-g** ITC confirms tight binding of all three enzymes to Scaf.4: SH3L-optiMHETase (**e**, K_D_ = 3.53 nM), FAST-PETase-CelK (**f**, K_D_ = 6.44 nM), and ICCG-ScaA (**g**, K_D_ = 5.25 nM). Data are presented as mean ± s.d. (n = 3). Statistical significance was determined by one-way ANOVA (ns: not significant, *p < 0.1, **p < 0.01, ***p < 0.001, ****p < 0.0001).

Degradation assays revealed several key insights (**Fig. 3b-d**). The result showed ICCG has slightly higher activity towards PET substrates compared to FAST at 30 °C. The inclusion of MHETase with ICCG led to near-complete conversion of MHET and BHET intermediates into TPA, and EG by implication. Moreover, combining ICCG, FAST, and MHETase greatly increased TPA productions to 33.07 mM (powder), 36.09 mM (film-L) and 0.93 mM (film-H). Scaf.4 further increased the efficiency to 44.66 mM (powder), 49.03 mM (film-L) and 1.32 mM (film-H). Finally, we confirmed the strong binding of all three enzymes (FAST, MHETase, and ICCG) to the scaffold, demonstrating stability of Complex-D (**Fig. 3e-g**). Overall, Complex-D further enhanced PET biodegradation efficiency and achieved high intermediate conversion.

### Improve SPEED performance through reaction optimization

To optimize the performance of the three-enzyme Complex-D and improve enzyme cost-effectiveness, we systematically investigated its PET degradation efficiency under various reaction conditions (**Fig. 4a**), beginning with the impact of enzyme-scaffold ratio (**Fig. 4b**). While increasing enzyme concentration could enhance product release (**Fig. 4b**, bar graph), ratios exceeding 0.5:1 (enzyme:Scaf.4) led to diminishing benefits from the scaffold (**Fig. 4b**, dot graph and **Supplementary Fig. 7**). This suggests that at higher concentrations, the scaffold becomes saturated, and the surplus of free enzyme does not contribute to the synergistic effect, reducing the overall cost-effectiveness. Consequently, we adopted the 0.5:1 enzyme-to-scaffold ratio for subsequent optimizations.

**Fig. 4:**
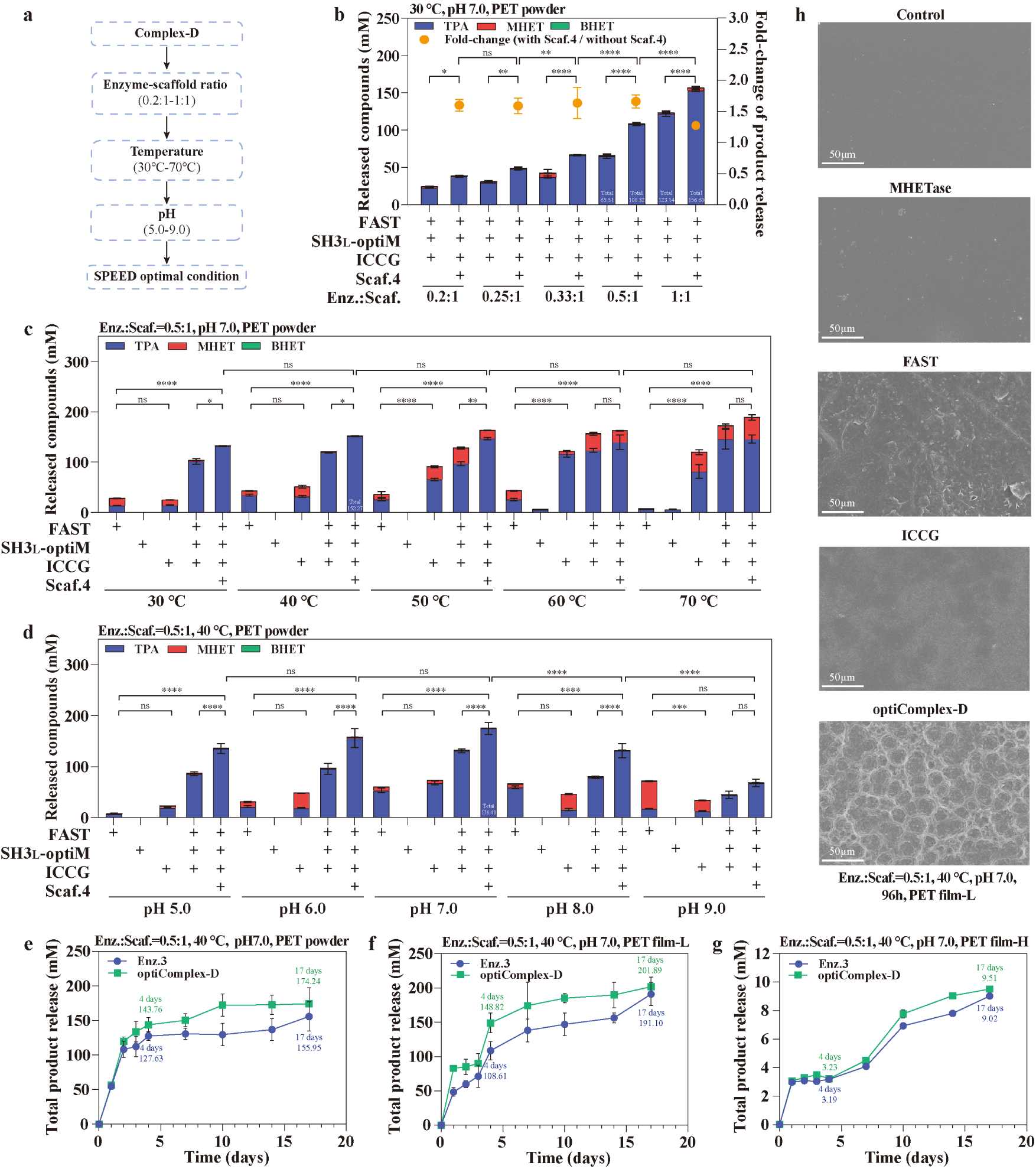
Reaction optimization enhances performance of the three-enzyme complex. **a** Strategy for optimizing Complex-D reaction conditions. **b** Effect of the enzyme:scaffold molar ratio on total product release from PET powder (bars, left axis) and the fold-change in activity conferred by the scaffold (dots, right axis). An optimal ratio of 0.5:1 was selected. Reactions contained 100 mg PET powder, enzymes with a constant 1 μM scaffold, were incubated at 30 °C, pH 7.0, for 96 h. **c** Influence of temperature on PET degradation efficiency and final products (0.5:1 enzyme:scaffold ratio, pH 7.0, 96 hours, 100 mg PET powder). **d** Impact of pH on PET degradation efficiency and final products (0.5:1 enzyme:scaffold ratio, 40 °C, 96 h, 100 mg PET powder). **e-g** Long-term (17-day) degradation time-course analysis for optiComplex-D vs. a non-scaffolded enzyme mixture (Enz.3) on PET powder (**e**), film-L (**f**), and film-H (**g**) under 40 °C, pH 7.0, with 0.5:1 enzyme:scaffold ratio. **h** Scanning electron microscopy (SEM) images of film-L after 96 hours of treatment at 40 °C, pH 7.0, with no enzyme (Control), single enzyme (MHETase, FAST, ICCG), and optiComplex-D. Data are presented as mean ± s.d. (n = 3). Statistical significance was determined by one-way ANOVA (ns: not significant, *p < 0.1, **p < 0.01, ***p < 0.001, ****p < 0.0001).

Next, we investigated the influence of temperature and pH on Complex-D performance (**Fig. 4c,d**). While previous findings indicate optimal temperatures for free FAST and ICCG are approximately 40 °C and 60 °C, respectively ^27^, our results for the complex showed that increasing the temperature above 40 °C led to an accumulation of intermediates and reduced TPA production, likely due to adverse effects on the less thermostable MHETase (**Supplementary Fig. 8**). Consequently, the highest total product yield (176.40 mM, with 174.71 mM as TPA) was achieved at 40 °C and pH 7.0, conditions consistent with the optimum for MHETase activity ^28^.

To directly assess the impact of scaffolding on enzyme kinetics, we compared the performance of the scaffold-binding enzymes against their free counterparts (**Supplementary Fig. 9**). These results showed that while the optimal reaction temperature and pH did not change upon scaffold binding, the scaffolded FAST and ICCG, but not MHETase, exhibited both higher PET degradation efficiency and increased tolerance to sub-optimal conditions, such as elevated pH and temperature.

In addition, the robustness of Complex-D under these optimal conditions was validated across diverse PET substrates, including PET powder, films of low and high crystallinity, and post-consumer commercial bottles (**Supplementary Fig. 10**). The system demonstrated significantly enhanced degradation activity across all tested materials over the controls, achieving conversion to the final monomer.

Finally, the catalytic stability and substrate versatility of the optimized Complex-D were further assessed through long-term PET degradation experiments (**Fig. 4e-g**). The complex exhibited robust and sustained hydrolysis across all PET substrates over 17 days. It is noteworthy that these optimized conditions, which balance the kinetics of the three-enzyme cascade, also allow robust performance of the free enzyme mixture, though its activity remained lower than that of the scaffolded complex. Scanning electron microscopy (SEM) confirmed this enhanced activity, revealing that treatment with the optimized Complex-D induced significantly greater surface erosion on PET film and a more pronounced reduction in PET powder particle size after 96 hours compared to the individual enzymes (**Fig. 4h** and **Supplementary Fig. 11**).

### Tailored MOF immobilization enhances catalytic stability and reusability

To improve the system’s stability and reusability for potential industrial translation, we explored enzyme immobilization strategies. We first immobilized optiComplex-D onto the metal-organic framework ZIF-8 ^29^, selected for its favorable properties including low cost and tunable properties ^30,31^. Because balancing enzyme stability and substrate accessibility is a critical in heterogeneous biocatalysis, so we prepared three distinct configurations: (i) surface immobilization, (ii) incomplete encapsulation, and (iii) full encapsulation (**Fig. 5a** and **Supplementary Fig. 12**).

**Fig. 5:**
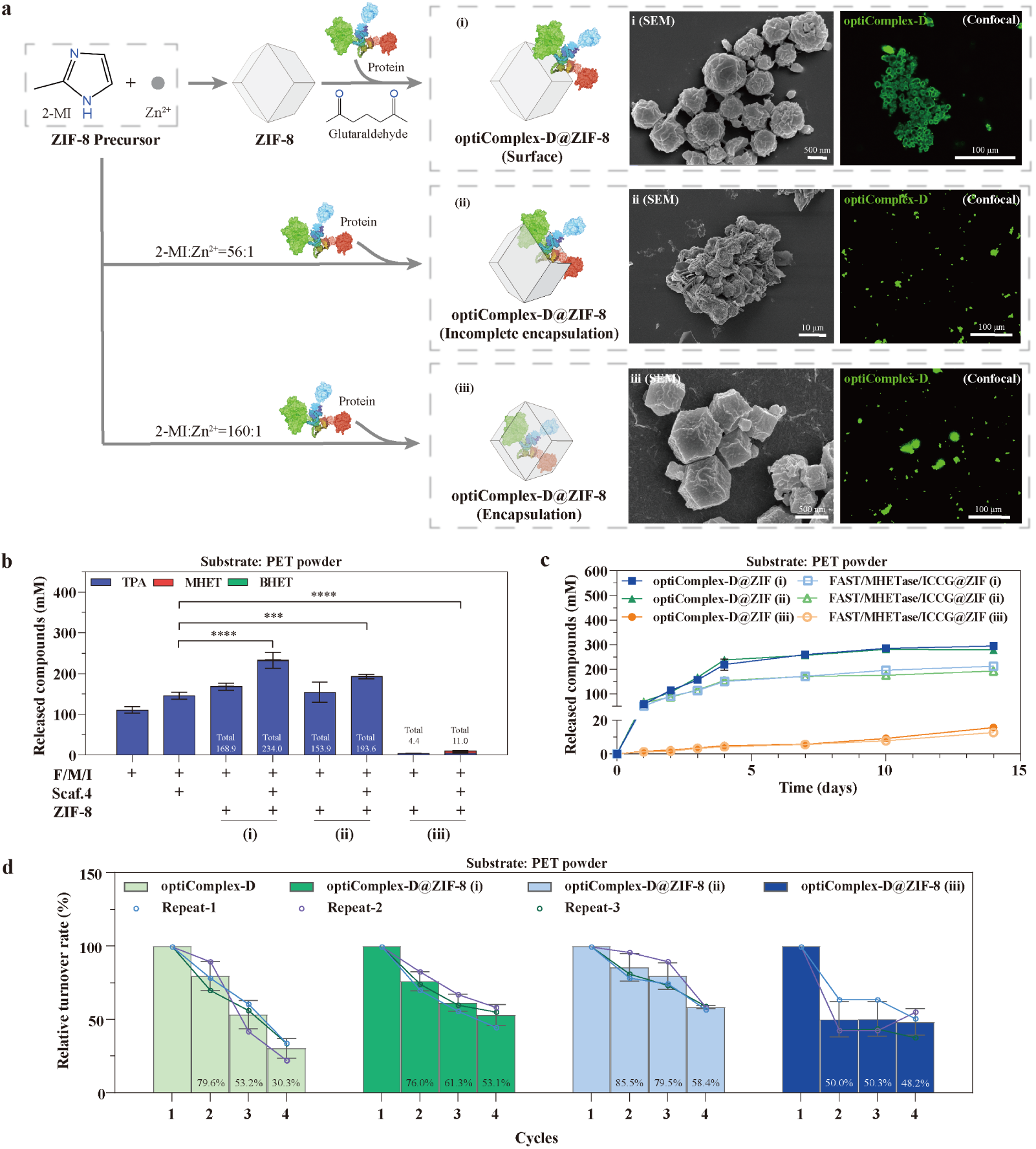
ZIF-8 immobilization enhances complex stability and reusability. **a** Schematic of three ZIF-8 immobilization strategies: (i) surface immobilization via glutaraldehyde crosslinking, (ii) incomplete encapsulation using a 2-MI:Zn²⁺ ratio of 56:1, and (iii) full encapsulation using a 160:1 ratio, following an established protocol ^34^. Successful immobilization and the distinct morphology of each configuration were confirmed by SEM and fluorescence microscopy. 2-MI, 2-methylimidazole. Alexa Fluor® 488 conjugated anti-His antibody was used to detect His-tagged proteins. The loading efficiency of optiComplex-D@ZIF-8 was 75.26 % (i), 85.95 % (ii), 91.13 % (iii) respectively. **b** Total product yield after 96 h on PET powder comparing three immobilization configurations. **c** Long-term (14-day) activity of scaffolded (optiComplex-D) and non-scaffolded (F/M/I) enzymes immobilized in the three configurations. **d** Reusability of free vs. immobilized optiComplex-D over four 4-day cycles. Reactions were performed at 40 °C, pH 7.0, with 100 mg PET powder, 0.25 nmol enzymes in a 0.5:1 enzyme:scaffold ratio. Data are presented as mean ± s.d. (n = 3). Statistical significance was determined by one-way ANOVA (ns: not significant, *p < 0.1, **p < 0.01, ***p < 0.001, ****p < 0.0001).

As expected, catalytic activity was highly dependent on substrate access. The fully encapsulated complex (iii) showed minimal PET degradation, while the incompletely encapsulated (ii) and surface-immobilized (i) complexes both demonstrated catalytic activity, with the former showing the highest performance (**Fig. 5b**). Furthermore, ZIF-8 immobilization significantly improved the complex’s stability across a wider pH and temperature range (**Supplementary Fig. 13**). The scaffold’s synergistic function remained crucial post-immobilization, with scaffolded complexes outperforming non-scaffolded enzyme mixtures in long-term assays (**Fig. 5c**). Reusability was also improved, as the top-performing complex retained over 53 % of its initial activity after four 4-day cycles, substantially outperforming the ∼ 30 % retained by the free complex (**Fig. 5d** and **Supplementary Fig. 14**).

To benchmark our MOF-based approach, we also immobilized the complex on calcium phosphate (CaP) nanocrystals, a versatile biomaterial that can be used for enzyme immobilization ^32,33^ (**Supplementary Fig. 15a**). While the initial 4-day product yield of optiComplex-D@CaP was comparable to that of the ZIF-8 system (**Supplementary Fig. 15b** vs. **Fig. 5c**), the CaP matrix proved less robust in metrics critical for practical application. For instance, the ZIF-8 system demonstrated enhanced reusability, retaining over 53 % of its activity after four cycles, compared to 40.9 % for the CaP-immobilized complex (**Fig. 5d** vs. **Supplementary Fig. S15c**). Furthermore, ZIF-8 offered a wider operational window. While the thermal stability of both systems was limited by the enzymes themselves, the ZIF-8 immobilized complex retained a greater proportion of its activity at 70 °C compared to the CaP system (**Supplementary Fig. 13a,c** vs. **14d**). The inherent chemical instability of CaP led to its dissolution and loss of activity at pH below 6.0 (**Supplementary Fig. 15e**), a limitation not observed with the ZIF-8 matrix. We also noted that the optiComplex-D@CaP formulation was unstable during cold storage, releasing the enzyme complex over time.

Together, these findings demonstrate that a tailored immobilization strategy using ZIF-8 preserves the catalytic advantages of the scaffolded architecture while conferring the enhanced stability and reusability beneficial for industrial translation.

### PET upcycling through scaffold-mediated biocatalysis

The modularity of the SPEED platform allows for the integration of PET depolymerization with downstream valorization pathways ^35–37^. To evaluate this capability, we engineered a second multi-enzyme assembly, Complex-E, designed to convert the PET degradation product ethylene glycol (EG) to glycolic acid (GA) ^38^, a versatile C2 platform chemical. GA has direct applications in dermatology and cosmetics and also serves as the monomer for poly(lactide-co-glycolide acid) (PLGA)-a high-performance biodegradable polymer used to create absorbable surgical sutures and drug delivery systems ^39–41^. This synthetic pathway employs the sequential action of 1,2-propanediol oxidoreductase (FucO) ^42^ and glycolaldehyde dehydrogenase (AldA) ^38^, with *Lactobacillus reuteri* NADH oxidase (LrNox) ^35^ regenerating the required NAD^+^ cofactor, a similar strategy building upon our prior work ^43^ (**Fig. 6a,b**).

**Fig. 6:**
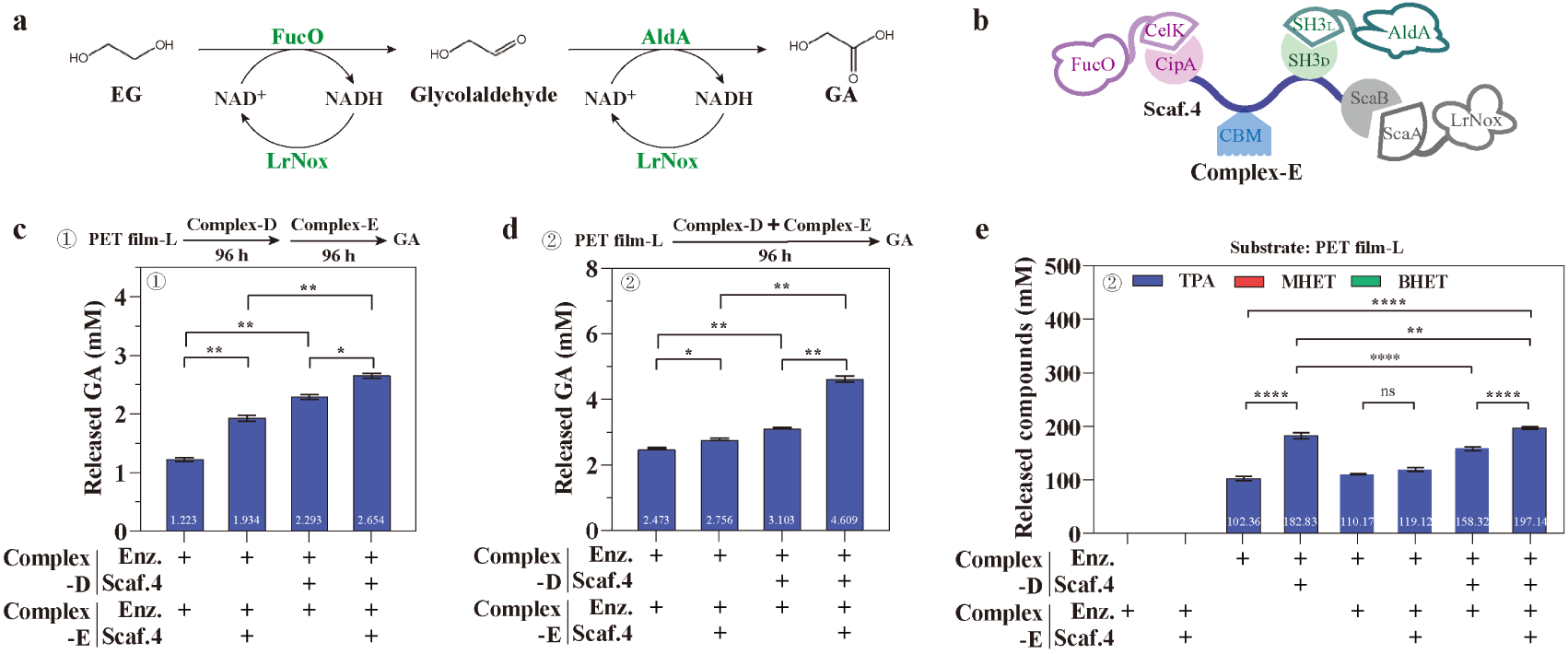
Modular scaffolds for one-pot PET depolymerization and monomer upcycling. **a** The upcycling cascade converts EG to glycolic acid (GA) via a three-enzyme pathway with NAD^+^ regeneration. **b** Schematic of the upcycling enzyme complex (Complex-E). **c** GA production in a sequential reaction, where Complex-E is added after a 96 h PET degradation step. **d** GA production in a simultaneous, one-pot reaction containing both Complex-D and Complex-E from the start, showing a 1.74-fold improvement over the sequential approach. **e** PET degradation performance under different enzyme and scaffold combinations. All reactions were performed at 40 °C, pH 7.0, for 96 hours, with 0.25 nmol of each individual enzyme and enzyme:scaffold molar ratios of 0.5:1 for optiComplex-D and 1:1 for Complex-E. Data are presented as mean ± s.d. (n = 3). Statistical significance was determined by one-way ANOVA (ns: not significant, *p < 0.1, **p < 0.01, ***p < 0.001, ****p < 0.0001).

We evaluated GA production by comparing sequential and simultaneous reaction schemes. A sequential approach, where Complex-E was added after a 96-hour PET degradation, yielded 2.65 mM GA (**Fig. 6c**). In contrast, a simultaneous, one-pot reaction with both complexes yielded 4.61 mM of GA in the same total time, a 1.74-fold increase (**Fig. 6d**). This enhanced performance likely stems from the continuous conversion of EG, improving the overall pathway flux. The importance of the scaffold in orchestrating this complex cascade was evident. When the upcycling enzymes (FucO, AldA, LrNox) were added as free proteins alongside optiComplex-D, PET degradation was significantly inhibited (**Fig. 6e**, compare bars 4 and 7). This result suggests that the free enzymes competed for binding sites on the Scaf.4 scaffold, disrupting the depolymerization cascade. Thus, assembling enzymes for each stage into distinct, scaffolded complexes is crucial for one-pot efficiency. This tandem reaction demonstrates the application of the SPEED design principle for engineering complex, multi-step biocatalytic processes.

### SPEED platform adapted for whole-cell biocatalysis

To demonstrate the SPEED platform’s potential for scalable biorecycling, we adapted it for a whole-cell biocatalyst system using yeast surface display, which avoids costly enzyme purification steps. In our system, an engineered yeast strain displays the cohesin scaffold on its cell surface, a second strain was engineered to secrete the dockerin-fused hydrolases (FAST-PETase and MHETase) into the medium (**Fig. 7a**). We selected the industrial yeast *Pichia pastoris* SMD1168 for its high-density growth and protein secretion capabilities ^44,45^ (**Supplementary Fig. 16**), and the scaffold was displayed to the cell surface using the SED1 anchor protein, which demonstrated the highest efficiency among several candidates (**Fig. 7b**). We also confirmed the high copy numbers of the expression cassettes in the engineered strains (**Supplementary Table 3**).

**Fig. 7:**
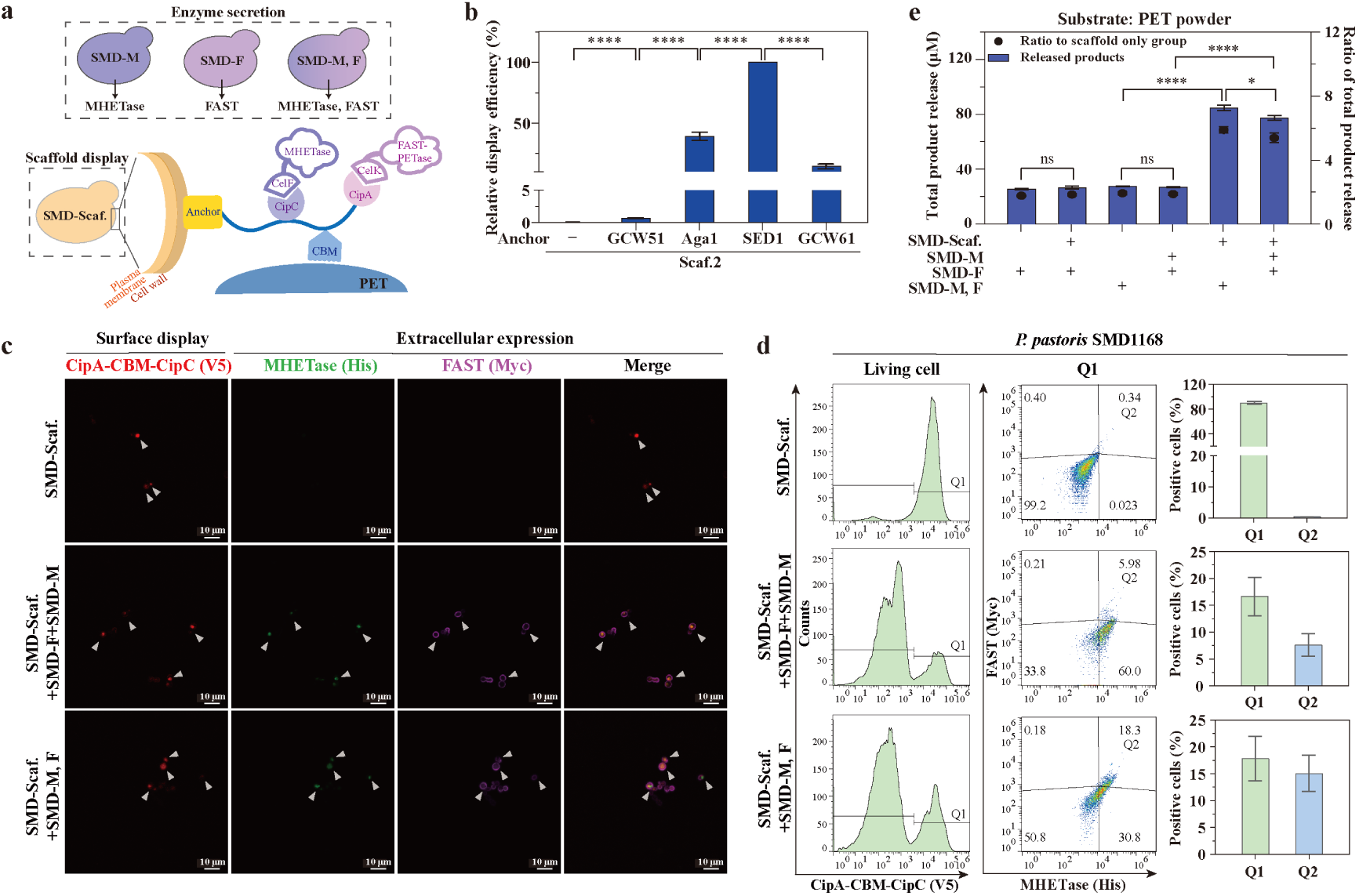
SPEED-enabled live cell system for PET degradation. **a** Design of the co-culture system in *P. pastoris*, where one strain displays the cohesin scaffold and a second strain secretes dockerin-fused enzymes for self-assembly on the cell surface. **b** Screening of anchor proteins for scaffold display, identifying SED1 as the most efficient. **c**, **d** Confocal microscopy (**c**) and flow cytometry (**d**) confirm the co-localization of secreted enzymes (MHETase-His, FAST-Myc) on the surface of scaffold-displaying yeast (Scaf.-V5). **e** PET degradation assays showing that the co-culture system with the scaffold-displayed complex significantly outperforms the control culture containing only free, secreted enzymes. Reactions used 100 mg PET powder and were incubated for 48 h at 30 °C, pH 7.0 after a 96-h protein expression induction. The ratio of enzyme-secreting strains to the scaffold-displaying strain was 10:1. Data are presented as mean ± s.d. (n = 3). Statistical significance was determined by one-way ANOVA (ns: not significant, *p < 0.1, **p < 0.01, ***p < 0.001, ****p < 0.0001).

Co-culturing these two strains resulted in the spontaneous, self-assembly of the enzyme complex onto the yeast cell surface, which was confirmed by confocal microscopy and flow cytometry (**Fig. 7c,d**) though we could not observe the individual enzyme on SDS-PAGE. We then compared the PET degradation efficiency of this whole-cell system against a control culture containing only the enzyme-secreting strain (without the scaffold-displaying yeast). The surface-displayed SPEED complex exhibited a 3.05-fold higher PET degradation activity than the secreted enzymes alone (**Fig. 7e**). This result demonstrates that the scaffold-mediated proximity and synergy are maintained in a complex *in vivo* environment, providing a promising proof-of-concept for developing robust, industrially relevant live cell systems for PET recycling.

## Discussion

In this study, we constructed a modular protein scaffold platform for PET biorecycling by assembling multi-enzyme cascades via orthogonal cohesin-dockerin and SH3 ligand/domain interactions. This spatial organization addresses the kinetic limitations observed in non-scaffolded enzyme mixtures by facilitating substrate channeling, thereby mitigating the accumulation of inhibitory intermediates (MHET, BHET) and increasing conversion to the final monomers (TPA, EG). We further evaluated the platform’s modularity through three proofs-of-concept directed toward potential industrial translation: immobilizing the complex on metal-organic frameworks to improve catalytic stability and enzyme reusability; extending the cascade to couple depolymerization with ethylene glycol upcycling; and adapting the assembly for a scalable live-cell system via yeast surface display (**Fig. 8**).

**Fig. 8:**
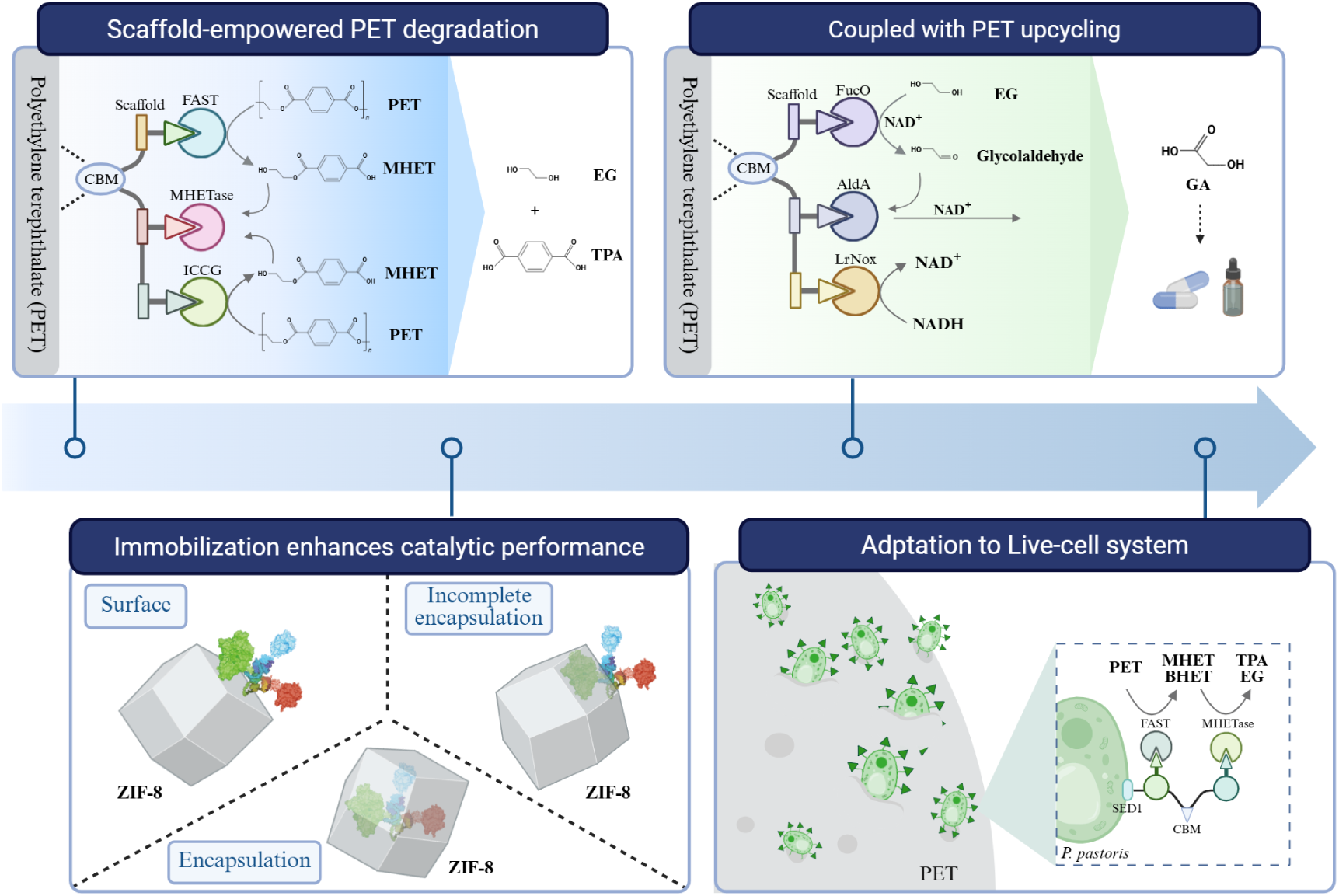
Schematic summary of the PET biorecycling system reported in this study. Top left: the scaffold-mediated PET degradation cascade. Bottom left: structural configurations for anchoring the enzyme complex onto metal-organic frameworks. Top right: coupling of the system to convert the ethylene glycol monomer into glycolic acid. Bottom right: assembly of the biocatalytic system via yeast surface display.

Building on previous multi-enzyme strategies, this scaffold-mediated assembly improved the product yield per mole of enzyme (**Supplementary Table 2**). Regarding substrate conversion extents (**Supplementary Table 4**), the optiComplex-D degraded 16.40% of high-crystallinity (40.82%) PET powder after four days. Immobilization on ZIF-8 increased this 4-day conversion to 27.14%. Over an extended 14-day reaction, the immobilized complexes achieved conversion extents of 33.49% for the high-crystallinity powder (40.82%) and 36.72% for a lower-crystallinity (17.99%) PET film.

These conversion extents reflect the use of untreated substrates, representing distinct experimental parameters from studies reporting near-complete depolymerization. For example, Tournier *et al*. utilized amorphized PET, higher enzyme loading, and a pH-stat bioreactor to chemically drive the reaction equilibrium ^13^. Similarly, Lu *et al*. evaluated post-consumer products with initial crystallinity ranging from 1.2% to 6.24% ^14^. Rather than maximizing bulk conversion via substrate pretreatment or continuous reactor optimization, this study evaluates normalized catalytic efficiency and targeted monomer production on untreated, high-crystallinity materials. These data indicate that structural scaffolds facilitate monomer production on PET substrates.

While this study describes a scaffold-mediated platform for PET biorecycling, further research is necessary for industrial translation. The current experiments were not evaluated under high-solids loading conditions, and techno-economic analyses are required to determine cost-effectiveness. Future development should focus on integrating thermally and chemically stable enzymes to broaden the operational parameters, as well as expanding the available upcycling pathways to establish viable biorefining routes. In summary, the modularity and scaffold-mediated spatial organization evaluated in this work provide a framework for constructing biocatalytic systems for plastic waste conversion.

## Methods

### Plasmid construction

Genes encoding enzymes and scaffolds were commercially synthesized (Tsingke, China), with all key primer sequences used in this study provided in **Supplementary Table 5.** Standard cloning techniques were employed using *E. coli* DH5α. Briefly, FAST-PETase (FAST), ICCG, FucO, AldA, LrNox and scaffold variants were cloned into pET28a vectors, while MHETase constructs utilized pET28a and pColdII vectors. Synthesized genes and pET28a vectors were digested with NdeI and XhoI (37 °C, 1h), followed by T4 DNA ligase-mediated ligation (25 ℃, 2h). This generated plasmids pET28a-FAST-PETase-CelK, pET28a-MHETase-CelF, pET28a-ICCG-ScaA, pET28a-FucO-CelF, pET28a-SH3_L_-AldA, pET28a-LrNox-ScaA, pET28a-CipA-CBM-CipC, pET28a-CipA-CBM-SH3_D_, and pET28a-CipA-CBM-SH3_D_-ScaB. In enzyme-scaffold complexes, enzymes were fused with native dockerin from cellulase enzyme CelK ^46^ and CelF ^47^ to construct FAST-PETase-CelK, FucO-CelF and MHETase-CelF, respectively. Meanwhile, the corresponding cohesin CipA and CipC were tethered together to form scaffold protein CipA-CipC (Scaf.1). Scaf.1 incorporates an additional carbohydrate-binding module (CBM) from *Clostridium thermocellum* ^23,48^, a module that enhances protein affinity for PET substrates ^49^, to form CipA-CBM-CipC (Scaf.2). The protein-protein interaction pair SH3_L_-SH3_D_ ^24^ and doc-coh pair ScaA-ScaB ^26^ were used to construct SH3_L_-AldA, CipA-CBM-SH3_D_ (Scaf.3), CipA-CBM-SH3_D_-ScaB (Scaf.4), SH3_L_-optiMHETase, ICCG-ScaA, LrNox-ScaA, respectively.

To construct the live-cell PET degradation system, *Pichia pastoris* SMD1168 was employed. The pPIC9K vector was utilized for the insertion and expression of MHETase, and Scaf.3. To engineer strain with exogenous gene insertion, primer pairs pScf-F/pScf-R were employed to amplify CipA-CBM-CipC-SED1, while primer pairs pF-F/pF-R and pM-F/pM-R were utilized to amplify FAST and MHETase, respectively. Additionally, primers pPIC9K-F/pPIC9K-R were used to amplify pPIC9K linearized vector. The pPICZC vector was utilized for the insertion of MHETase. MHETase and pPICZC linearized vector were amplified using primer pairs pZM-F/pZM-R and pPICZC-F/pPICZC-R. The linearized gene and vector were connected by seamless cloning to form the expression vectors pPICZC-FAST, pPIC9K-MHETase, pPIC9K-CipA-CBM-CipC-SED1, and pPICZC-MHETase. The expression vectors were linearized by SacI digestion and transformed into *P. pastoris* SMD1168. The transformants were cultured in histidine-deficient (SD-his) plates and screened for strains with high copies of the exogenous gene insertion by SD-his plates containing 5 mg/mL G418.

To reposition SH3_L_ from the C-terminus to the N-terminus of optiMHETase, sequential PCR was performed with primer pairs M1-F/M-R, M2-F/M-R, and M3-F/M-R. To remove SH3_L_ from SH3_L_-optiMHETase (as control), PCR was performed with primer pairs M1-F/Md-R. For C-terminal His-tag of MHETase, PCR was conducted using primers his-F/his-R. Subsequent digestion with NdeI and XbaI, followed by T4 DNA ligase-mediated recombination, generated plasmids pColdII-MHETase-CelF, pColdII-optiMHETase-CelF, pColdII-optiMHETase-SH3_L_, pColdII-SH3_L_-optiMHETase, and pColdII-optiMHETase. All protein constructs were designed with both N- and C-terminal His tags.

### Recombinant protein expression and purification

For Recombinant protein expression, *E. coli* C41 (DE3) cells were transformed with pColdII-based plasmids, while *E. coli* BL21 (DE3) and *E. coli* Rosetta (DE3) cells were used for pET28a constructs. Transformants were grown in LB media supplemented with appropriate antibiotics (100 µg/mL ampicillin for pColdII, 500 µg/mL kanamycin for pET28a) at 37 °C until OD_600_ reached 0.6-0.7. IPTG (0.5 mM or 1.0 mM final concentration for MHETases, 1.0 mM for others) induced expression overnight (at 16 °C or 37 ℃ for MHETases, 16 ℃ for others). Cells were harvested by centrifugation and stored at −80 °C.

Cells were resuspended in lysis buffer (50 mM sodium phosphate, 300 mM NaCl, pH 8.0), disrupted by sonication, and the supernatant was collected by centrifugation. HisTrap HP column purification employed an ÄKTA pure system (Cytiva, China). Bound protein was washed with 85 % binding buffer (20 mM sodium phosphate, 30 mM imidazole, 500 mM NaCl, pH 7.4) with 15 % elution buffer (20 mM sodium phosphate, 500 mM imidazole, 500 mM NaCl, pH 7.4) and eluted with a linear gradient of 15-100 % elution buffer. Eluates were concentrated (Amicon Ultra, Merck Millipore, China), buffer-exchanged to 50 mM sodium phosphate (SP buffer, pH 7.0), and assessed for purity/concentration by NanoDrop™ One/One^C^ (Thermo Fisher, U.S.A.). The high purity of all proteins was confirmed by SDS-PAGE (**Supplementary Fig. S1**), and representative FPLC purification profiles are shown in **Supplementary Fig. S2**. Proteins were stored in storage buffer (50 % glycerol in SP buffer, pH 7.0) at −80 °C.

### Binding assay of protein scaffolds to PET

Following previously established protocols ^49^, PET powder (crystalline, Xing-xiang new materials, China) or PET film (amorphous, Goodfellow, UK) were washed with 10 % SDS, water, and SP buffer (pH 7.0). CipA-CipC (Scaf.1) or CipA-CBM-CipC (Scaf.2) were incubated at 2.5, 5, and 7.5 µM with 100 mg PET powder in 200 µL SP buffer (pH 7.0) overnight at 4 °C. Samples were centrifuged (12000 g, 1 min), washed twice with SP buffer (pH 7.0), and residual protein in solution was quantified using a BCA Protein Quantification Kit (Yeason, China). Binding protein concentration was calculated as follows (C, concentration; before, before incubation with PET; after, after incubation with PET):

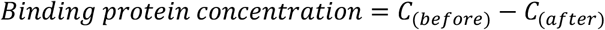

### Binding assay of enzymes to scaffolds

Enzyme-scaffold protein interactions were analyzed by MicroCal PEAQ-ITC Automated (Malvern Panalytical, Netherland). Enzymes were placed in the reaction cell, and scaffold protein titrations were performed at 25 °C with 18 injections of 2 µL substrate ^50^. Heats generated by dilution were corrected, and data were analyzed using the MicroCal PEAQ-ITC Analysis Software (one set of sites model).

For live-cell system, flow cytometry was used to analyze enzyme-scaffold protein interactions. After a 96-hour induction at 30 °C, the yeast cells were collected by centrifugation at 4000 g for 10 min at 4 °C, then washed and diluted with DPBS. Subsequently, 1×10^7^ cells and 0.1 μL of each antibody (V5 tag antibody, Invitrogen 451098; His tag antibody, Cell Signaling Technology 14930; or Myc tag antibody, R&D Systems IC3696P) were incubated for 45 min, washed and detected through flow cytometry.

### Characterization of crystallinity of PET substrates

Thermal crystallinity of PET substates was determined by differential scanning calorimetry (DSC 8000, PerkinElmer) (**Supplementary Fig. 3**). PET substrates (75 µm powder; low crystallinity PET film, film-L; high crystallinity PET film, film-H; Nongfu Spring water bottle, NFbottle) were heated from 30 ℃ to 300 ℃ at a heating rate of 30 ℃/min, kept at 300 ℃ for 1 min, and then cooled to 0 ℃ at 5 min with same rate of 30 ℃/min ^14,49^. The theoretical heat of melting value (ΔHm°) of 100 % crystallized PET was 140.1 J/g. Percentage crystallinity was calculated with the heat of melting (ΔHm) and cold crystallization (ΔHc) values, as the following equation:

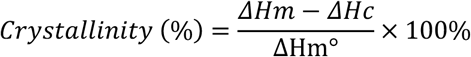

### Synthesis and characterization of MOF-immobilized enzyme complexes

For synthesis of FAST/MHETase/ICCG/Scaffold@ZIF-8 (optiComplex-D@ZIF-8) and FAST/MHETase/ICCG@ZIF-8, a certain amount of proteins (0.25 nmol individual enzyme of FAST, MHETase and ICCG with/without scaffold to form enzyme complexes), an aqueous solution of 2-methylimidazole (790 mM, 1.75 mL for incompletely encapsulated crystals, 5 mL for completely encapsulated crystals) and 0.25 mL Zn(NO_3_)_2_·6H_2_O (97.5 mM) were stirred at 500 g for 5 min. To synthesize ZIF-8, 1.75 mL 2-MI (790 mM) and 0.25 mL Zn(NO_3_)_2_·6H_2_O (97.5 mM) was stirred. Subsequently, the mixture was left undisturbed for 3 h at 4 °C ^34^. For synthesis of enzyme complexes immobilization on ZIF-8 carrier, 20 mg prepared ZIF-8 with a certain amount of proteins was cultured in 5 mL SP buffer containing 50 mM glutaraldehyde at 4 ℃ for 3 h ^51^. Finally, the prepared Enz.Scaf@ZIF-8, Enz@ZIF-8 and ZIF-8 were collected by centrifugation at 3500 g for 10 minutes and dispersed into 0.5 mL SP buffer.

For characterizations, confocal microscopy images were recorded by Nikon confocal laser scanning microscope; SEM (ZEISS GeminiSEM 300) were used to measure the immobilized samples.

### Synthesis and characterization of CaP-immobilized enzyme complexes

The FAST/MHETase/ICCG@CaP and FAST/MHETase/ICCG/Scaffold@CaP (optiComplex-D@CaP) were synthesized according to the previously reported method with slight modification by mixing solution I with solution II ^52,53^. For solution I, 1 mg Ca_3_(PO_4_)_2_ was added to 2 mL of 50 mM sodium phosphate buffer with different pH. For solution II, 0.25 nmol individral enzymes with/without 0.5 nmol scaffold (Scaf.4) were added to 2 mL of 100 mM CaCl_2_ solution. Solution I and II were balanced for 1 h at 4 ℃, respectively, then mixed with equal volume. After incubation for 12 h at 4 ℃, the precipitate was washed thrice and dispersed into 0.5 mL SP buffer to obtain optiComplex-D@CaP and FAST/MHETase/ICCG@CaP. For characterizations, confocal microscopy images were recorded by Nikon confocal laser scanning microscope.

### Loading efficiencies of enzymes in ZIF-8 and CaP

The loading efficiencies of compatible enzymes in ZIF-8 and CaP were evaluated with BCA Protein Quantification Kit (Yeason, China). Specifically, collecting the supernatant from the washing process and final step synthesis of enzyme@ZIF-8 or enzyme@CaP, then detecting the concentration and volume of the supernatant. The loading efficiencies were then determined as the ratio of embedded enzymes amount to the initial amount of enzymes ^34^, which was calculated based on the following equation (m_(before)_, the initially added enzyme amount (mg); C_(after)_, the enzyme concentration in the supernatant (mg/mL); V_(after)_, the volume of supernatant (mL)):

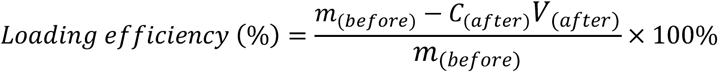

### Exogenous gene insertion analysis

The insertion copy number of exogenous genes was determined by quantitative real-time PCR (qPCR), following an established protocol ^44^. The genomic DNA of engineered *P. pastoris* SMD1168 was extracted by the Yeast Genomic DNA Extraction Kit (TIANGEN, DP307). Genomic DNA of *P. pastoris* SMD1168_His4::CipA-CBM-CipC-SED1_, *P. pastoris* SMD1168_His4::FAST_, *P. pastoris* SMD1168_His4::MHETase_ and *P. pastoris* SMD1168_His4::FAST-MHETase_ were used as templates, respectively, for the amplification of the genes CipA-CBM-CipC, FAST, MHETase with the primer pairs qC-F/qC-R, qF-F/qF-R, and qM-F/qM-R; the housekeeping gene GAPDH was amplified with the primer pairs qGAPDH-F/qGAPDH-R to obtain the copy number of the exogenous protein inserted in the engineered strains. All primer sequences were listed in **Supplementary Table 3**.

### Establishment and characterization of live-cell PET biodegradation system

For live-cell systems, surface-displaying and secretory-expressing engineered yeast strains were individually pre-cultured in BMGY (1 % yeast extract, 2 % peptone, 100 mM potassium phosphate buffer, 1.34 % amino acid-free yeast nitrogen source, and 1 % (v/v) glycerol, pH 6.0) at 30 °C for 12 h, after which the number of cells was measured in the pre-cultured products. *P. pastoris* SMD1168_His4::enzyme_ was mixed with *P. pastoris* SMD1168_His4::scaffold_ in a 10:1 ratio, resulting in a final system comprising 2×10^7^ *P. pastoris* SMD1168_His4::scaffold_. The expression medium BMMY (1% yeast extract, 2 % peptone, 100 mM potassium phosphate buffer, 1.34 % amino acid-free yeast nitrogen, and 1 % (v/v) methanol, pH 6.0) was employed for protein expression and enzyme-scaffold assembly. Expression was conducted at 30 °C, with the medium augmented with 0.5 % methanol at 24-hour intervals over a period of 96 hours.

Confocal microscopy images and flow cytometry assays were recorded by Nikon confocal laser scanning microscope. His Tag Alexa Fluor® 488-conjugated Antibody (IC0501G, R&D Systems), V5 Tag Monoclonal Antibody (451098, Invitrogen), Human c-Myc PE-conjugated Antibody (IC3696P, R&D Systems) to detect MHETase and/or FAST with His tag, FAST with Myc tag, scaffold with V5 tag, respectively.

For surface displaying assays mediated by different anchors, displaying efficiency was calculated with median florescence intensity (MFI) and copy number by flow cytometry and qPCR, as the following equation:

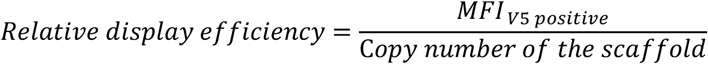

### Degradation assay of PET plastics

Standard reactions contained 100 mg PET (powder, low or high crystallinity PET films), 0.2 μM individual enzyme, 500 µL reaction buffer (50 mM sodium phosphate, 0.5 mM CaCl_2_, 10 % DMSO, pH 7.0). For complex assembly, enzymes (0.2 μM) were pre-mixed with scaffold (1 µM) for 40 min at 25 ℃. Unless otherwise stated, degradations proceeded for 96 h at 30 °C.

To optimize the enzyme:scaffold ratio within SPEED, ratios ranged from 0.2:1 to 1:1 (constant scaffold protein 1 μM). Effects of pH (5.0-9.0, 30 °C), temperature (30-70 °C, pH 7.0), and PET substrate type (PET powder, low and high crystallinity PET films, commercial bottle) were also analyzed.

For PET upcycling reaction, PET degrading enzymes (FAST, MHETase, ICCG) were combined with Scaf.4 (1 μM, CipA-CBM-SH3_D_-ScaB) in an optimized ratio. Similarly, the upcycling enzymes (FucO, AldA, LrNox), which facilitate the synthesis of valuable products from PET, were mixed with Scaf.4 (0.5 mM) in a molar ratio of 1:1. The mixed proteins were incubated at 25 ℃ for 40 minutes to achieve complex assembly. Following assembly, the upcycling complex was incorporated into the PET degradation product, which was reacted for four days under optimal conditions. Alternatively, the PET degradation complex was mixed with the upcycling complex and 100 mg PET Film-L for further conversion. The reactions were conducted in 500 µL systems for 2 to 4 days in SP buffer (with 1 mM additional NAD^+^).

For live-cell PET degradation, a mixture of 1×10^9^ induced *Pichia pastoris* SMD1168 was collected, then placed in 400 μL of sterile glycine-NaOH buffer (pH 7.0) containing 100 mg PET powder to facilitate the degradation reaction. The degradation reaction was conducted at 30 °C for 48 h. All reactions were terminated at 95 °C for 10 min, supernatants were 0.22 µm filtered, and released products were quantified by HPLC (High Performance Liquid Chromatography).

### MHETase degradation assay

To assess the intermediates degradation ability of MHETase, 0.5 μM SH3_L_-optiMHETase was combined with 5 mM MHET and BHET in 500 µL reaction buffer (50 mM sodium phosphate, 0.5 mM CaCl_2_, 10 % DMSO, pH 7.0). The reactions were constructed at varying temperatures (30 °C, 40 °C, 50 °C, 60 °C, 70 °C) for 96 h and terminated by heating to 95 °C for 10 min. Supernatants were 0.22 µm filtered, and released products were quantified by HPLC.

### HPLC analysis

TPA, MHET, BHET and GA, standards were obtained from Aladdin (China). The analysis of the samples was conducted using an Agilent 1260 LC system equipped with G1315A diode array detector (DAD) and ZORBAX StableBond C18 column (4.6×150 mm, 5 µm, Agilent) for TPA, MHET and BHET detection, as well as a refractive index detector (RID) and a Xtimate® Sugar-H column (3.8×300 mm, 5 µm, Welch Materials) for GA detection.

For TPA, MHET and BHET detection, the mobile phase consisted of 0.1 % formic acid (A) and acetonitrile (B); separation used a flow rate of 1 mL/min (80 % A / 20 % B, 10 min); detection was performed at 240 nm, 25 ℃ with C18 column ^25^.

For GA detection, the mobile phase consisted of 5 mM H_2_SO_4_ (Solvent as ddH_2_O); separation used a flow rate of 1 mL/min (15 min); detection was performed at 80 ℃ with Sugar-H column. Analyte concentrations were determined using calibration curves (10-500 mM). Samples were adaptively diluted with ultrapure water prior to analysis.

### Scanning electron microscopy (SEM)

Low-crystallinity PET film (6 mm diameter) and PET powder samples were analyzed by SEM (HuaSuan Technolygy, China) after degradation (40 °C, 96 h, pH 7.0). Samples were washed (10 % SDS, water, ethanol), dried, gold-coated, and imaged.

### Scaffold-enzyme structural prediction

In this study, three enzymes structures were obtained from the Protein Data Bank (https://www.rcsb.org/. FAST-PETase: 6IJ6, MHETase: 6QZ4, ICCG: 8JMP, CelK: 3P6B, SH3_L_: 1PRM, ScaA: 5N5P). The structure of the scaffold was predicted by AlphaFold2. After running the standard AlphaFold2 MSA generation pipeline against UniRef, MGnify, PDB, and BFD, the sequence with highest pLDDT greater than 0.9 is chosen as the final model prediction. The depiction of enzyme-scaffold complex was visualized by BioRender (https://www.biorender.com/).

### Statistical analysis

One-way analysis of variance (ANOVA) followed by normality test was employed to evaluate the significance of differences among different treatments. Unless specify, total products were used for the significance test. Statistical analysis were conducted using GraphPad Prism 9.0 software. ns: not significant, *p < 0.1, **p < 0.01, ***p < 0.001, ****p < 0.0001.

## Data Availability

All data are available in the main text, supplementary information and **Source Data**. Additional information related to this study is available from the corresponding author upon reasonable request.

## Acknowledgements

This study was supported by funding from the Shenzhen Bay Laboratory Startup Fund, the Shenzhen Outstanding Talents Training Fund, the National Natural Science Foundation of China (Grant Numbers: 32471546), Shenzhen Research Funding for Postdocs and the Key Research and Development Program of Shaanxi Province, China (Grant Numbers: 2022NY-055 and 2019NY-187). We acknowledge the intellectual and experimental contributions of all authors for this study and its manuscript preparation. We are particularly grateful for the insights gained from discussions with lab members in the Zhuobin Liang and Yanbin Lin labs. We would like to acknowledge Ziwei Luo for her technical assistance in the illustration of Figure. 1. We also extend our gratitude to the Haizhen Long and Jianchuan Wang lab at Shenzhen Bay Laboratory for their assistance in experiment setup. Finally, we thank the Research Core Facility of Shenzhen Bay Laboratory for their technical support, especially for the use of ÄKTA and HPLC instruments.

## Contributions

**Yujia Zhang:** Methodology, Investigation, Data Curation, Formal Analysis, Visualization, Writing – Original Draft. **Chongsen Li:** Investigation, Data Curation, Validation. **Ehsan Hashimi**: Investigation, Data Curation, Validation. **Enting Xu:** Investigation, Validation. **Xuemei Yang:** Methodology, Supervision, **Yanbing Lin:** Resources, Supervision. **Hui Gao:** Conceptualization, Methodology, Investigation, Data Curation, Formal Analysis, Writing – Original Draft, Funding acquisition. **Zhuobin Liang:** Conceptualization, Methodology, Supervision, Writing – Original Draft, Project administration, Funding acquisition.

## Extended Data

Source Data

## Supplementary information

### Supplementary Tables

**Supplementary Table 1.**
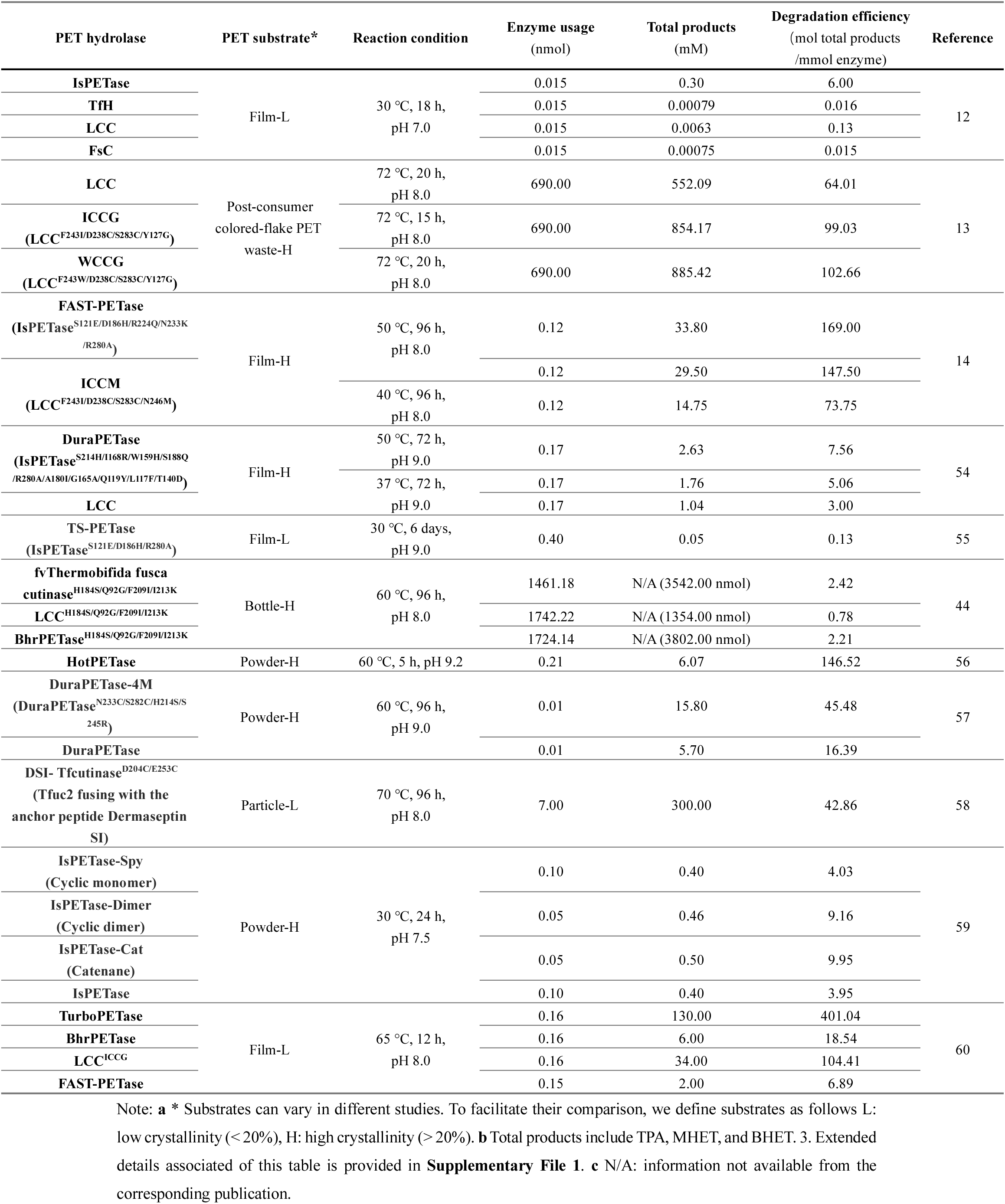
Key degradation parameters and efficiency of major PET hydrolases.

**Supplementary Table 2.**
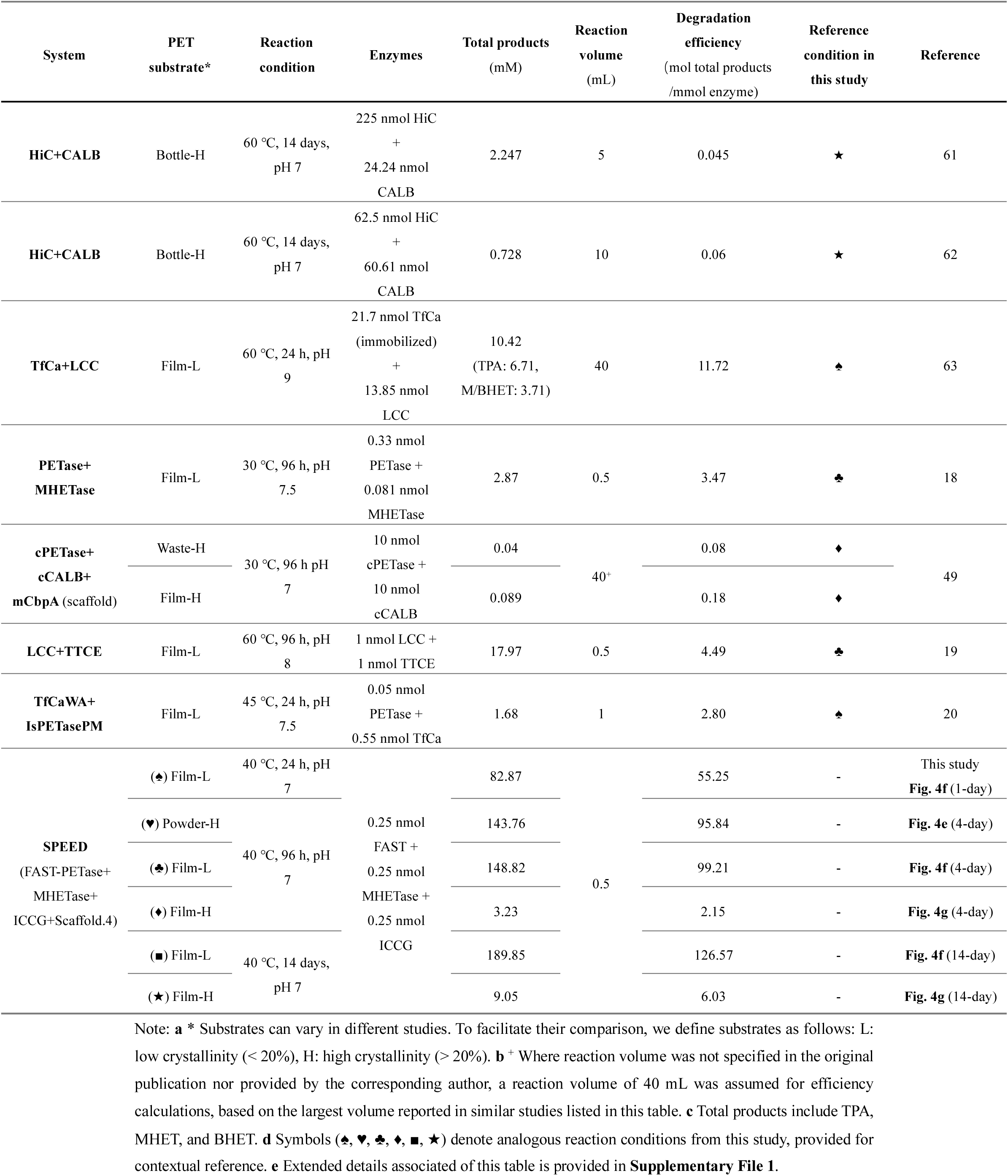
Current multi-enzyme systems for biodegradation of PET plastics.

**Supplementary Table 3.**
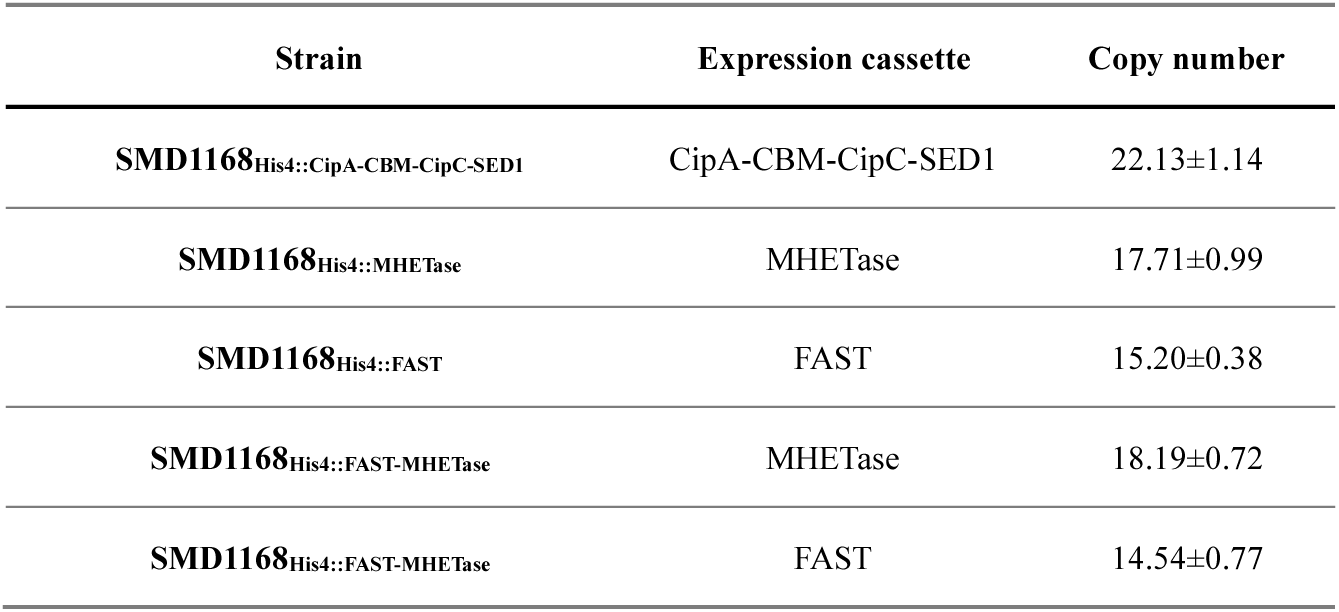
Copy number of enzyme and scaffold expression cassettes in the genome of the engineered *P. pastoris* strains.

| Strain | Expression cassette | Copy number |
| --- | --- | --- |
| **SMD1168**<sub>His4::CipA-CBM-CipC-SED1</sub> | CipA-CBM-CipC-SED1 | 22.13±1.14 |
| **SMD1168**<sub>His4::MHETase</sub> | MHETase | 17.71±0.99 |
| **SMD1168**<sub>His4::FAST</sub> | FAST | 15.20±0.38 |
| **SMD1168**<sub>His4::FAST-MHETase</sub> | MHETase | 18.19±0.72 |
| **SMD1168**<sub>His4::FAST-MHETase</sub> | FAST | 14.54±0.77 |

**Supplementary Table 4.** Calculated PET conversion extents for key reactions in this study.

| Reaction | Released<br>TPA<br>(mM) | Released<br>MHET<br>(mM) | Released<br>BHET<br>(mM) | Conversion<br>extent (% of<br>total substrate) | Reaction<br>condition | Data source |
| --- | --- | --- | --- | --- | --- | --- |
| **Complex-B**<br>**+PET powder** | 19.49 | 2.62 | 0.00 | 2.50% | 30 °C, pH 7.0,<br>4 days | Fig.1f, the last bar |
| **Complex-B**<br>**+PET Film-L** | 17.29 | 0.00 | 0.00 | 1.97% | 30 °C, pH 7.0,<br>4 days | Fig.1g, the last bar |
| **Complex-B**<br>**+PET Film-H** | 0.71 | 0.01 | 0.01 | 0.08% | 30 °C, pH 7.0,<br>4 days | Fig.1h, the last bar |
| **Complex-C**<br>**+PET powder** | 22.37 | 1.63 | 0.20 | 2.75% | 30 °C, pH 7.0,<br>4 days | Fig.2f, the last bar |
| **Complex-C**<br>**+PET Film-L** | 23.22 | 0.00 | 0.27 | 2.68% | 30 °C, pH 7.0,<br>4 days | Fig.2g, the last bar |
| **optiComplex-D**<br>**+PET powder** | 145.95 | 43.36 | 0.79 | 21.29% | 70 °C, pH 7.0,<br>4 days | Fig.4c, the last bar |
| **optiComplex-D**<br>**+PET powder** | 140.30 | 1.42 | 2.04 | 16.40% | 40 °C, pH 7.0,<br>4 days | Fig.4e, 4-day |
| **optiComplex-D**<br>**+PET powder** | 170.18 | 1.92 | 2.14 | 19.87% | 40 °C, pH 7.0,<br>17 days | Fig.4e, 17-day |
| **optiComplex-D**<br>**+PET Film-L** | 146.75 | 0.43 | 1.65 | 16.98% | 40 °C, pH 7.0,<br>4 days | Fig.4f, 4-day |
| **optiComplex-D**<br>**+PET Film-L** | 201.89 | 0.00 | 0.00 | 23.02% | 40 °C, pH 7.0,<br>17 days | Fig.4f, 17day |
| **optiComplex-D @ZIF-8(i)**<br>**+PET powder** | 215.73 | 3.74 | 0.00 | 24.99% | 40 °C, pH 7.0,<br>4 days | Fig.5c, 4 days'<br>point |
| **optiComplex-D @ZIF-8(i)**<br>**+PET powder** | 293.38 | 0.47 | 0.00 | 33.49% | 40 °C, pH 7.0,<br>14 days | Fig.5c, 14-day |
| **optiComplex-D @ZIF-8(ii)**<br>**+PET powder** | 237.19 | 0.63 | 0.23 | 27.14% | 40 °C, pH 7.0,<br>4 days | Fig.5c, 4-day |
| **optiComplex-D @ZIF-8(ii)**<br>**+PET powder** | 276.94 | 1.65 | 0.00 | 31.74% | 40 °C, pH 7.0,<br>14 days | Fig.5c, 14-day |
| **optiComplex-D @CaP**<br>**+PET Film-L** | 255.09 | 0.00 | 0.00 | 29.08% | 40 °C, pH 7.0,<br>4 days | Fig.S15b, 4-day |
| **optiComplex-D @CaP**<br>**+PET Film-L** | 319.84 | 2.44 | 0.00 | 36.72% | 40 °C, pH 7.0,<br>14 days | Fig.S15b, 14-day |
| **optiComplex-D +Complex-E**<br>**+PET powder** | 197.14 | 0.00 | 0.00 | 22.47% | 40 °C, pH 7.0,<br>4 days | Fig.6e, the last bar |

Note: PET conversion extent (%) was calculated from the total moles of released products (TPA, MHET, and BHET), as quantified by HPLC, relative to the initial molar quantity of PET repeating units (details in **Supplementary Methods**). The crystallinity of the substrates was determined to be 40.82% (PET powder), 17.99% (Film-L).

**Supplementary Table 5.**
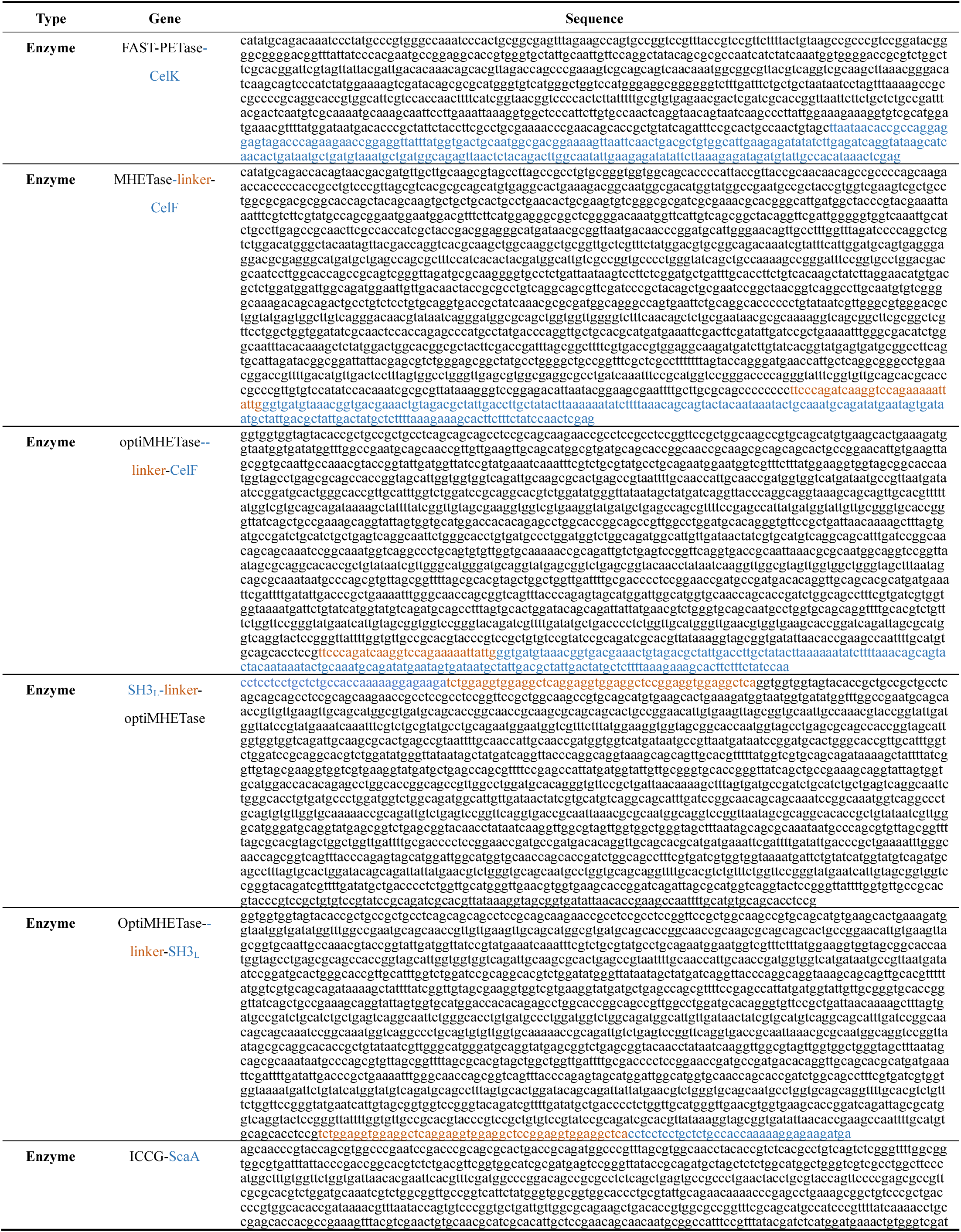
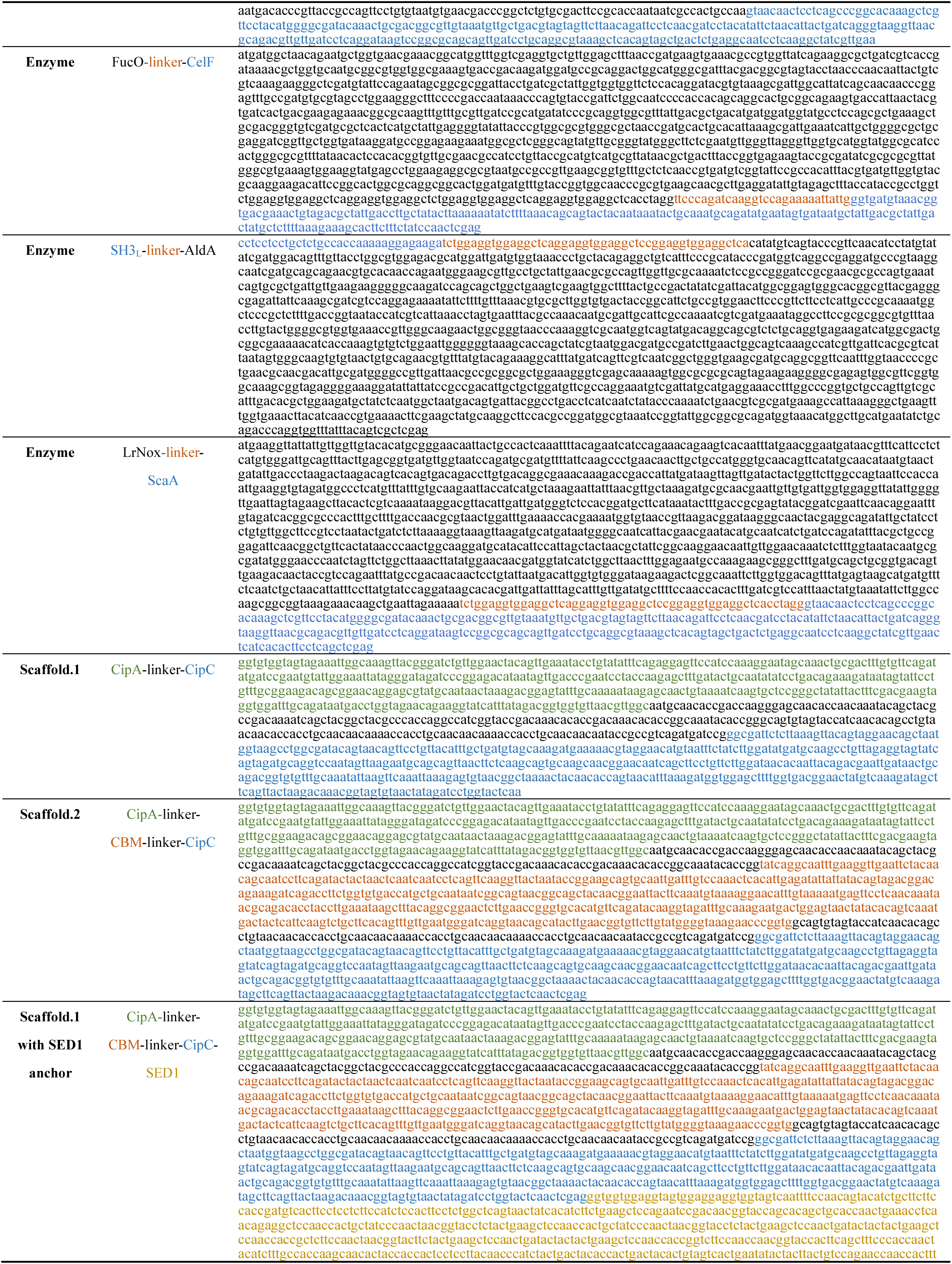
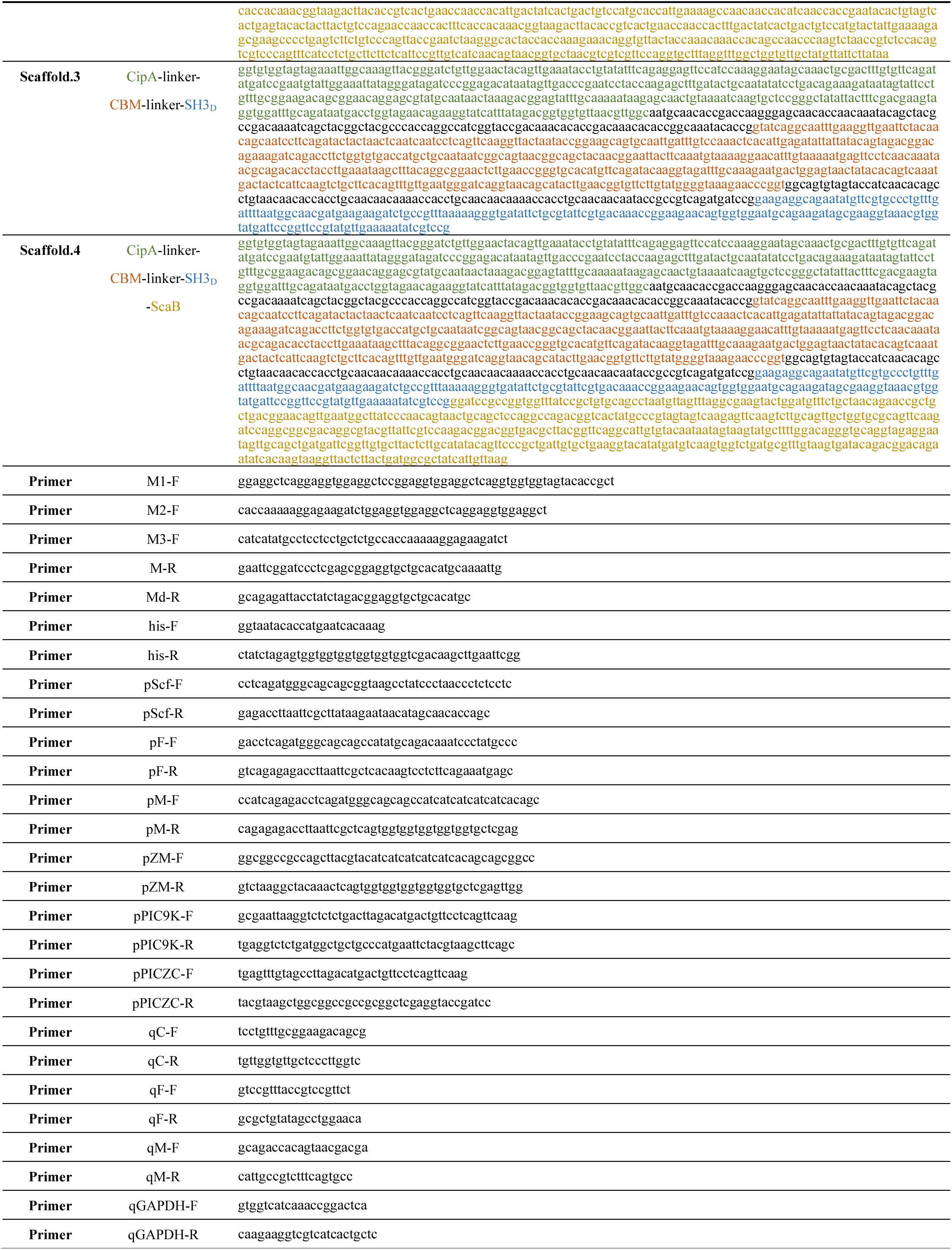
Information of key genes and primers of this study.

### Supplementary Figure

**Supplementary Fig. 1:**
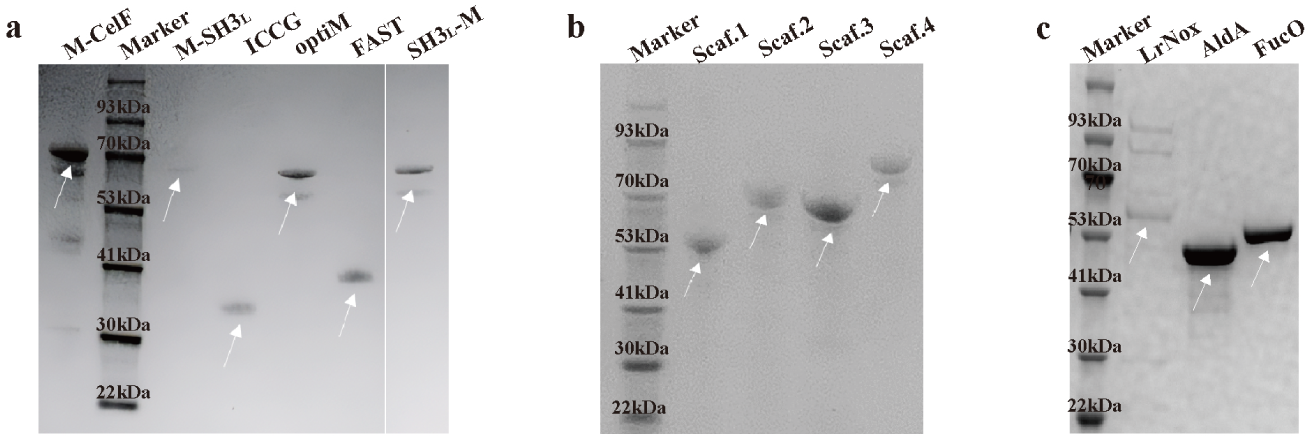
SDS-PAGE of purified recombinant enzymes and protein scaffolds used in this study. SDS-PAGE analysis of purified proteins stained with Coomassie Blue. **a** M-CelF: MHETase-CelF; optiM: codon optimized MHETase; M-SH3_L_: optiMHETase-SH3_L_; SH3_L_-M: SH3_L_-optiMHETase; ICCG: ICCG-ScaA; FAST: FAST-PETase-CelF. **b** Scaf.1: CipA-CipC; Scaf.2: CipA-CBM-CipC; Scaf.3: CipA-CBM-SH3_D_; Scaf.4: CipA-CBM-SH3_D_-ScaB. **c** LrNox: LrNox-ScaA; FucO: FucO-CelF; AldA: SH3_L_-AldA.

**Supplementary Fig. 2:**
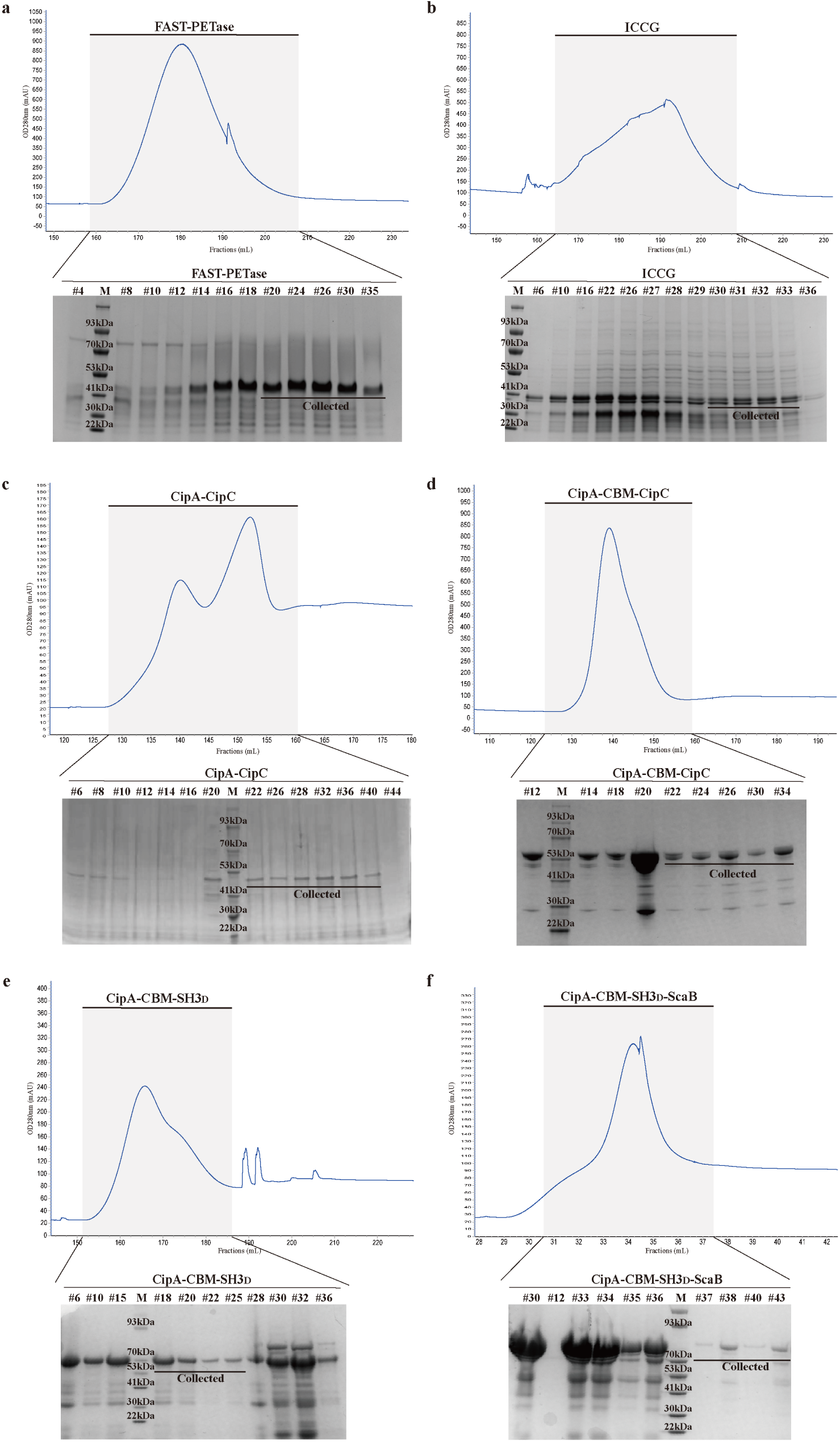
Purification profiles of key enzymes and scaffolds. **a-f** FPLC chromatograms (top) and corresponding SDS-PAGE analysis of elution fractions (bottom) for His-tag purified proteins. Shaded regions on chromatograms indicate the elution fractions from the HisTrap column subjected to SDS-PAGE analysis. **a** FAST-PETase-CelK. **b** ICCG-ScaA. **c** CipA-CipC (Scaf.1). **d** CipA-CBM-CipC (Scaf.2). **e** CipA-CBM-SH3_D_ (Scaf.3). **f** CipA-CBM-SH3_D_-ScaB (Scaf.4). M, protein molecular weight marker.

**Supplementary Fig. 3:**
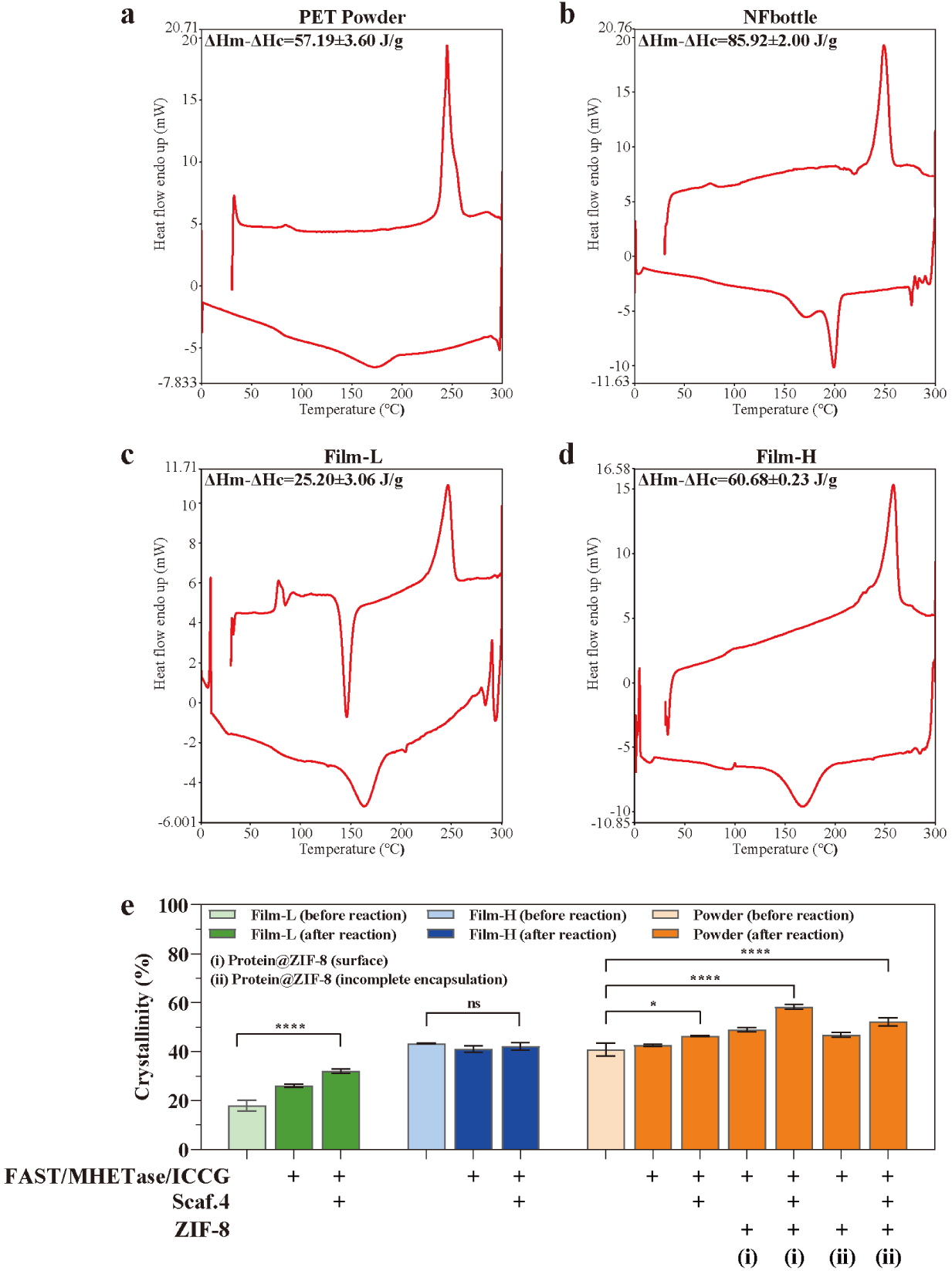
Crystallinity characterization of PET substrates. Differential scanning calorimetry (DSC) thermograms used to determine the crystallinity of PET substrates used in this study. **a** PET powder (40.82%). **b** Commercial water bottle (NFbottle, 61.33%). **c** Film-L (17.99%). **d** Film-H (43.31%). ΔHm, heat of melting; ΔHc, heat of cold crystallization. **e** Comparison of substrate crystallinity before and after a 96-h reaction with optiComplex-D or immobilized complexes at 40 °C, pH 7.0, using 100 mg PET substrate, 0.25 nmol individual enzymes, with an enzyme:scaffold molar ratio of 0.5:1. The apparent increase in crystallinity post-reaction is a known phenomenon; it occurs because the enzymes preferentially degrade the more accessible amorphous regions, thereby enriching the proportion of crystalline domains in the remaining material. Data are presented as mean ± s.d. (n = 3). Statistical significance was determined by one-way ANOVA (ns: not significant, *p < 0.1, **p < 0.01, ***p < 0.001, ****p < 0.0001).

**Supplementary Fig. 4:**
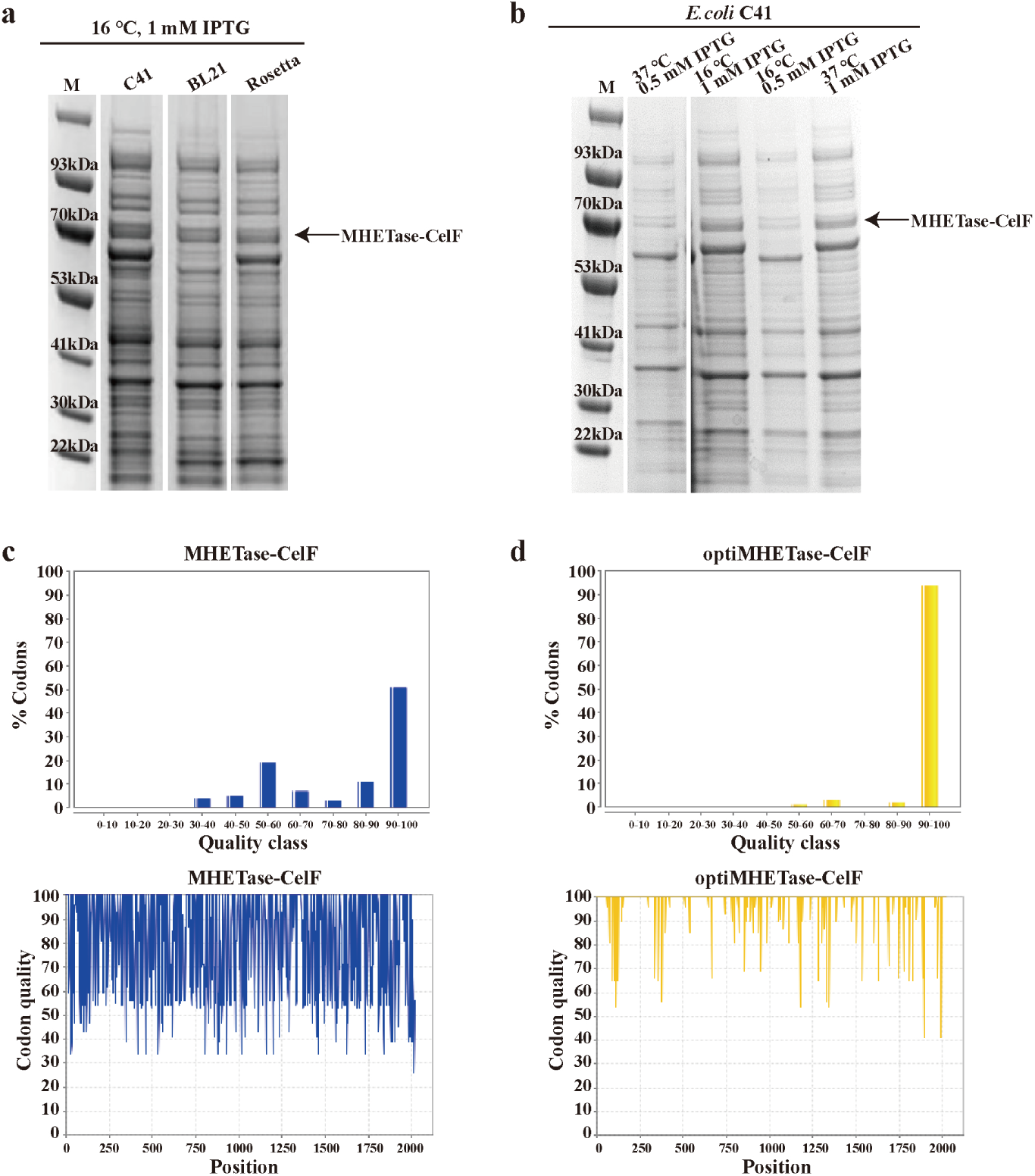
Optimization of MHETase expression in *E. coli*. **a** Screening of different *E. coli* expression strains, identifying C41(DE3) as the most effective host. **b** Optimization of induction temperature and isopropyl β-D-1-thiogalactopyranoside (IPTG) concentration in the C41(DE3) strain. Optimal expression was achieved at 16 °C with 1 mM IPTG for 18 h. M, protein molecular weight marker. The optimized expression condition was confirmed using *E. coli* C41 (DE3) for 12 hours at 16℃, with the addition of 1 mM IPTG for the induction. **c**, **d** Codon usage analysis before (**c**) and after (**d**) codon optimization for *E. coli* expression. The optimized gene shows a significantly improved distribution of high-quality codons (90-100 quality class). Results were calculated by GeneArt™ GeneOptimizer™ (Thermo Fisher).

**Supplementary Fig. 5:**
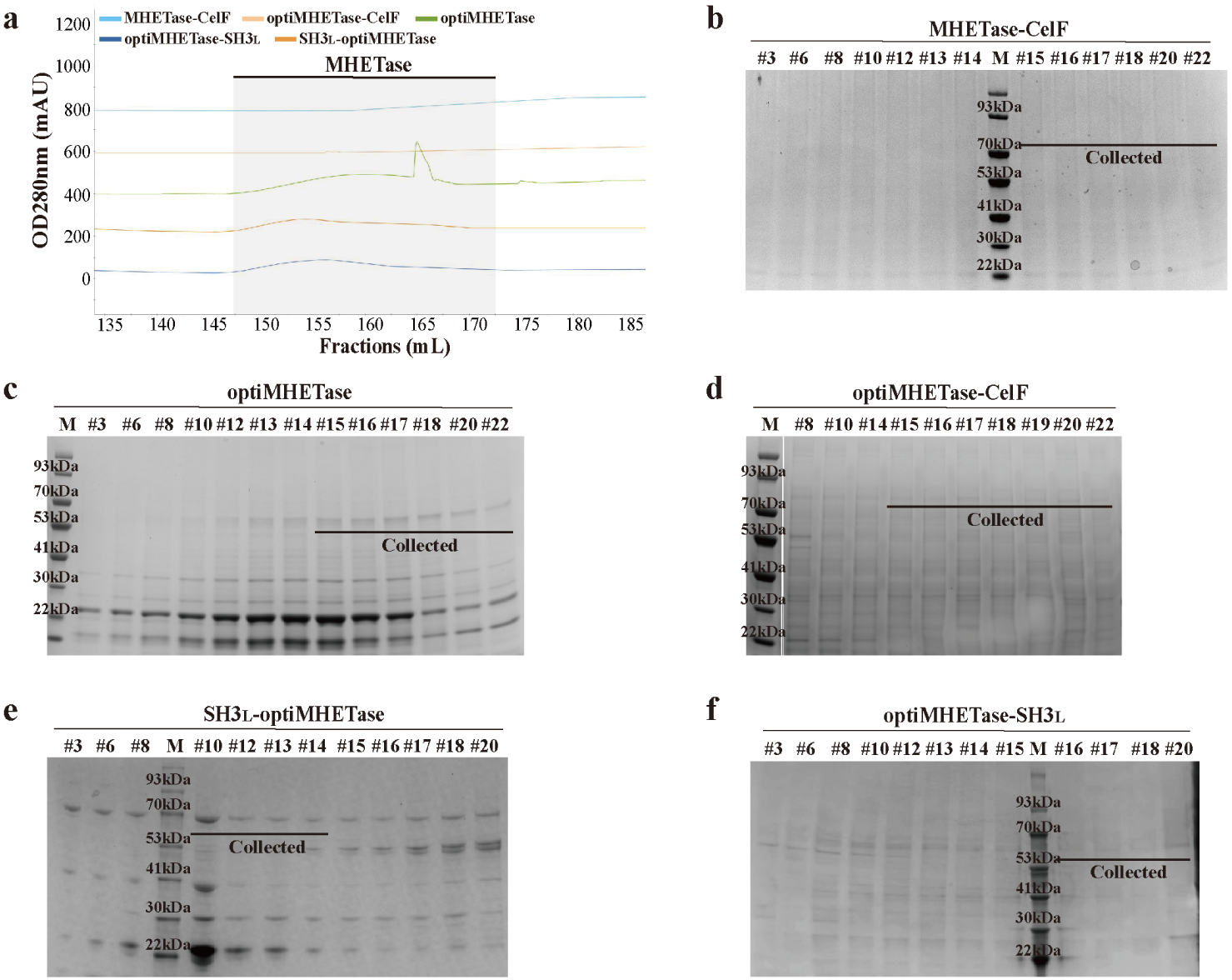
Purification profiles for MHETase variants. **a** Overlay of FPLC chromatograms for different MHETase constructs purified via His-tag affinity chromatography. **b-f** SDS-PAGE analysis of elution fractions for each construct, confirming the purity of the collected fractions (indicated by black bars).

**Supplementary Fig. 6:**
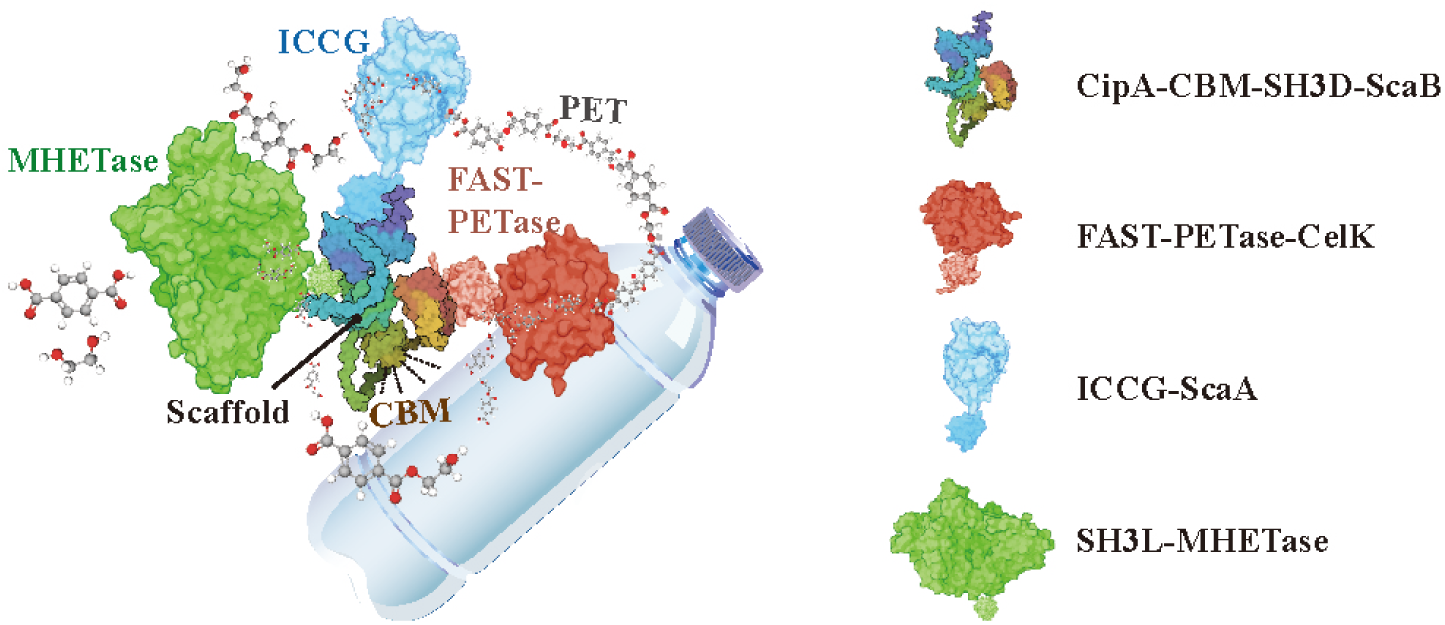
Structural model of the three-enzyme Complex-D. A model of the fully assembled complex, generated by integrating experimentally determined structures of the enzymes with an AlphaFold2-predicted structure of the Scaf.4 scaffold. Individual components are shown on the right for clarity.

**Supplementary Fig. 7:**
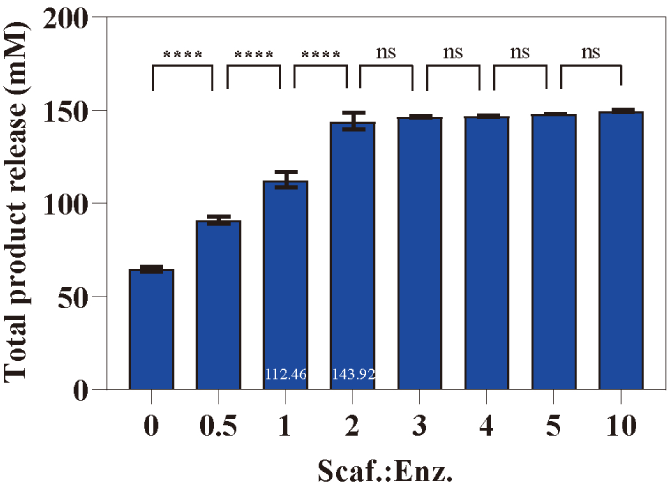
Effect of enzyme-to-scaffold molar ratio on Complex-D activity. PET degradation assays were performed with a fixed concentration of enzymes (0.25 nmol each of FAST, optiMHETase, ICCG) and varying concentrations of Scaf.4. The data show that activity plateaus at a scaffold:enzyme molar ratio of approximately 2:1, indicating saturation of the enzymes with scaffold. Reactions used 100 mg PET powder at 30 °C, pH 7.0, for 96 h. Data are presented as mean ± s.d. (n = 3). Statistical significance was determined by one-way ANOVA (ns: not significant, *p < 0.1, **p < 0.01, ***p < 0.001, ****p < 0.0001).

**Supplementary Fig. 8:**
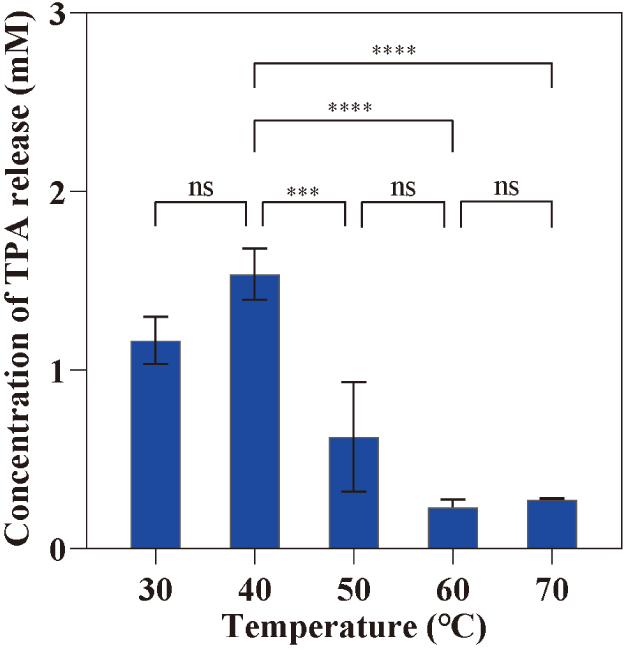
Effect of temperature on SH3_L_-optiMHETase activity. Activity of SH3L-optiMHETase was assayed on a mixture of 5 mM MHET and BHET at different temperatures with 0.5 μM enzyme at pH 7.0 for 96 h. The enzyme shows the highest activity for converting intermediates to TPA at 40 °C, consistent with the optimal temperature determined for the complete Complex-D cascade. The elevated TPA release observed in the 70 °C group was likely attributed to the self-degradation of the monomers under high temperatures. Data are presented as mean ± s.d. (n = 3). Statistical significance was determined by one-way ANOVA (ns: not significant, *p < 0.1, **p < 0.01, ***p < 0.001, ****p < 0.0001).

**Supplementary Fig. 9:**
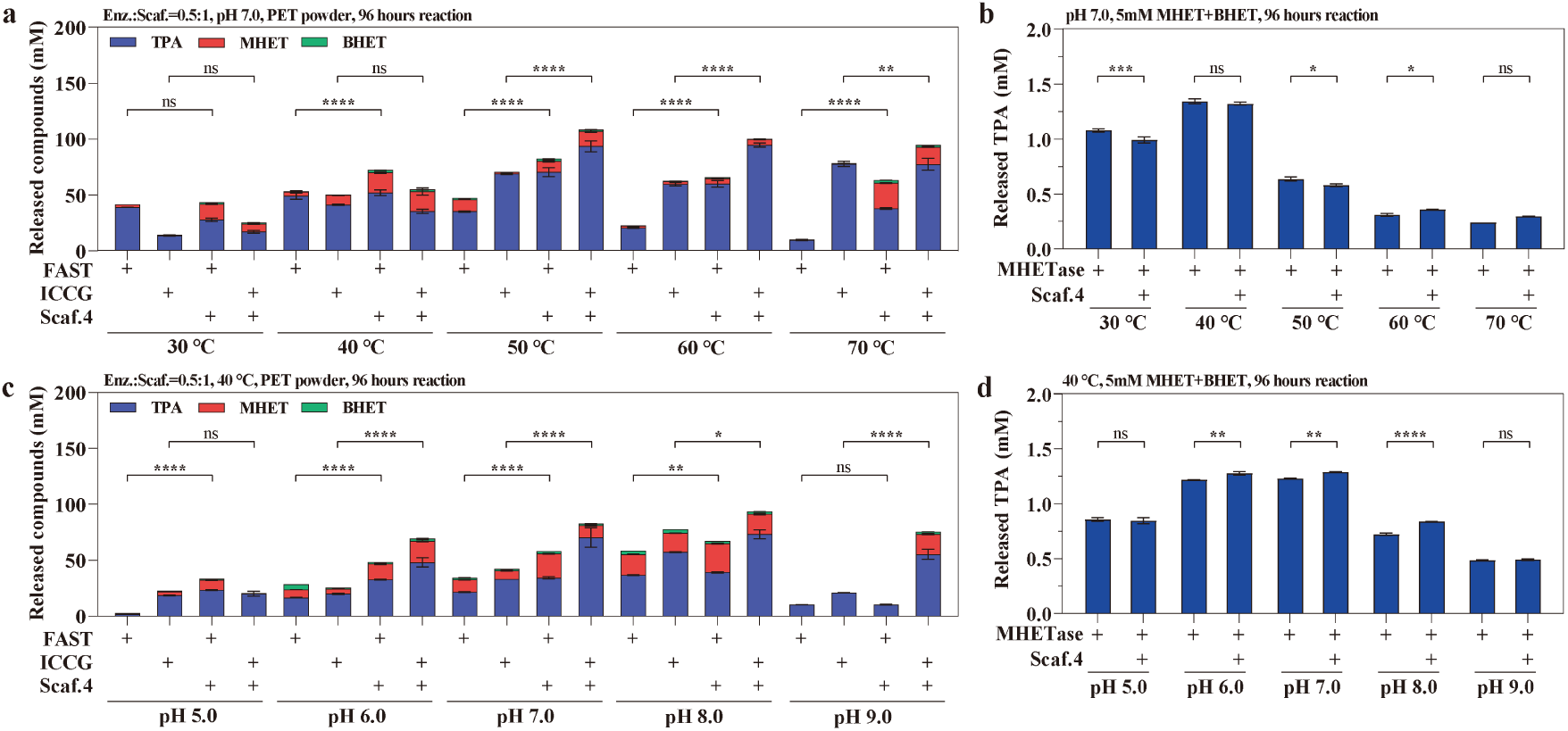
Comparison of scaffolded and free enzyme performance across varying pH and temperature conditions. Degradation of PET powder or intermediates MHET and BHET by the scaffolded enzymes was compared to an equimolar mixture of the corresponding free enzymes. **a-b** Temperature profile showing total product release across a range of temperatures (30-70°C) at pH 7.0. **c-d** pH profile showing total product release across a range of pH values (5.0-9.0) at 40°C. All reactions were performed for 96 hours using 100 mg PET powder or 5 mM MHET and BHET, using 0.25 nmol individual enzymes. Data are presented as mean ± s.d. (n = 3). Statistical significance was determined by one-way ANOVA (ns: not significant, *p < 0.1, **p < 0.01, ***p < 0.001, ****p < 0.0001).

**Supplementary Fig. 10:**
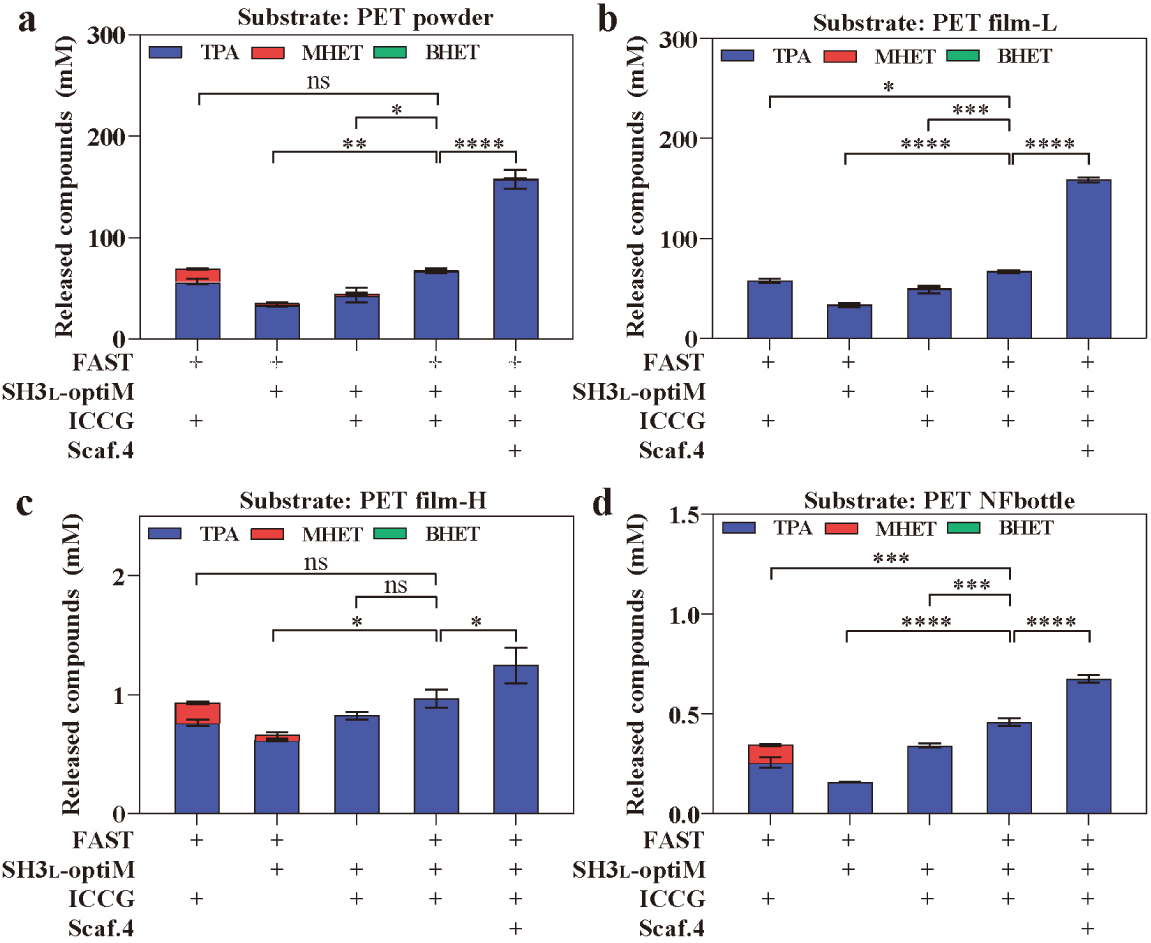
Performance of optiComplex-D across diverse PET substrates. Degradation assays under optimal reaction conditions (40 °C, pH 7.0, 96 h) comparing non-scaffolded enzyme mixtures with the fully assembled optiComplex-D. The scaffolded complex consistently shows superior performance on **a**, PET powder, **b**, low-crystallinity film (film-L), **c**, high-crystallinity film (film-H), and **d**, a commercial water bottle (NFbottle). All reactions contain 100 mg PET substrate, 0.25 nmol individual enzymes, with an enzyme:scaffold molar ratio of 0.5:1. Data are presented as mean ± s.d. (n = 3). Statistical significance was determined by one-way ANOVA (ns: not significant, *p < 0.1, **p < 0.01, ***p < 0.001, ****p < 0.0001).

**Supplementary Fig. 11:**
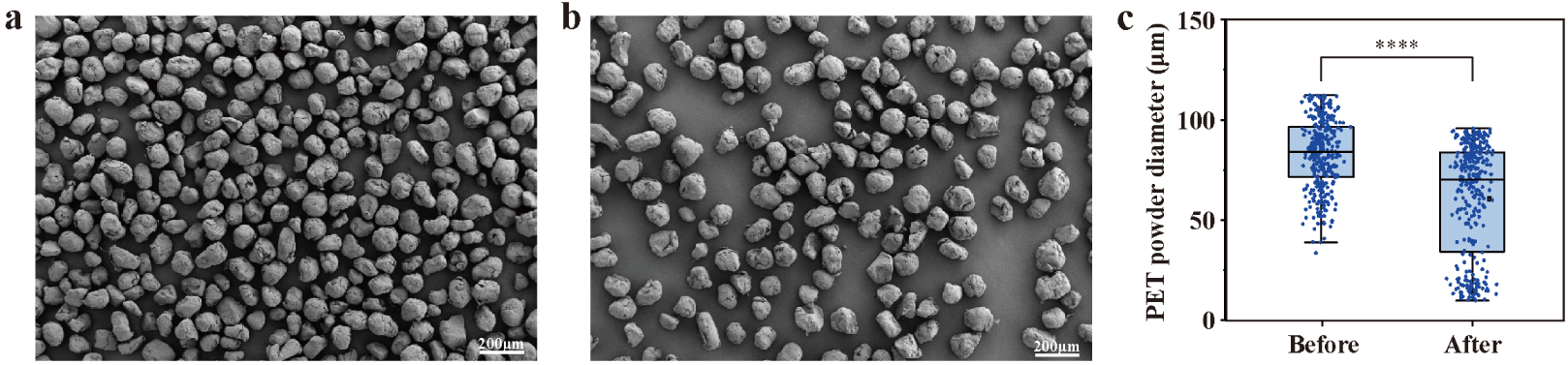
Enzymatic treatment reduces PET powder particle size. **a**, **b** SEM images of PET powder before (**a**) and after (**b**) a 96 h treatment with optiComplex-D at 40 °C, pH 7.0. **c** Boxplot analysis quantifying the significant decrease in particle diameter after enzymatic degradation. The box extends from the 25th to 75th percentiles, the line indicates the median, and whiskers show the min to max range. Statistical significance was determined by one-way ANOVA (****p < 0.0001).

**Supplementary Fig. 12:**
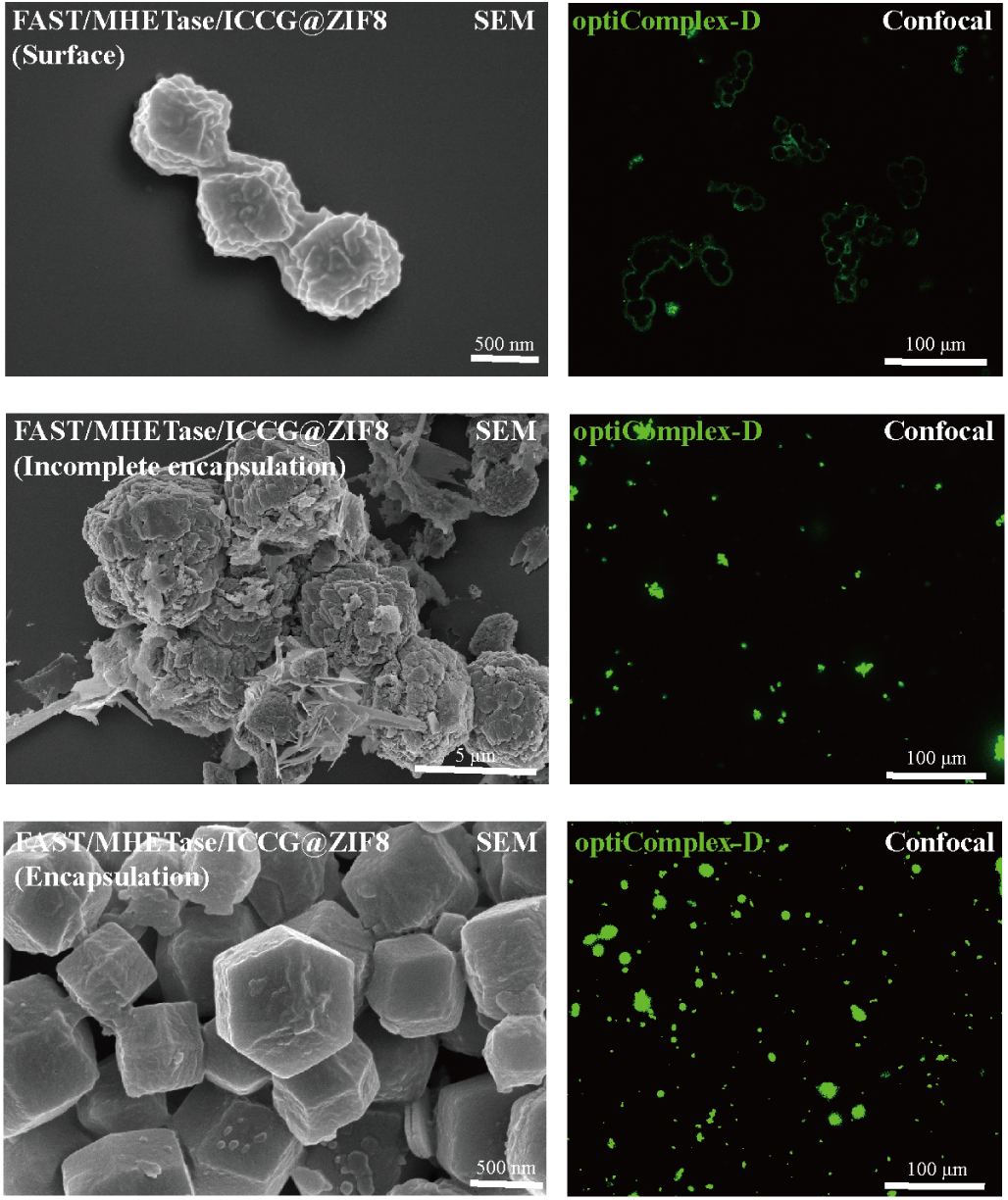
Characterization of non-scaffolded enzymes immobilized on ZIF-8. SEM and confocal microscopy images of a mixture of free enzymes (FAST-PETase, MHETase, and ICCG) immobilized on ZIF-8 using the three different strategies (surface, incomplete encapsulation, encapsulation), corresponding to the controls used in Fig. 5. His-tagged proteins were detected with an Alexa Fluor® 488 conjugated anti-His antibody. The loading efficiency of FAST/MHETase/ICCG@ZIF-8 was 78.11% (surface), 93.99% (incomplete encapsulation), and 90.14% (encapsulation). Confocal images serve as qualitative confirmation of successful immobilization; variations in fluorescence intensity are not for quantitative comparison of enzyme loading.

**Supplementary Fig. 13:**
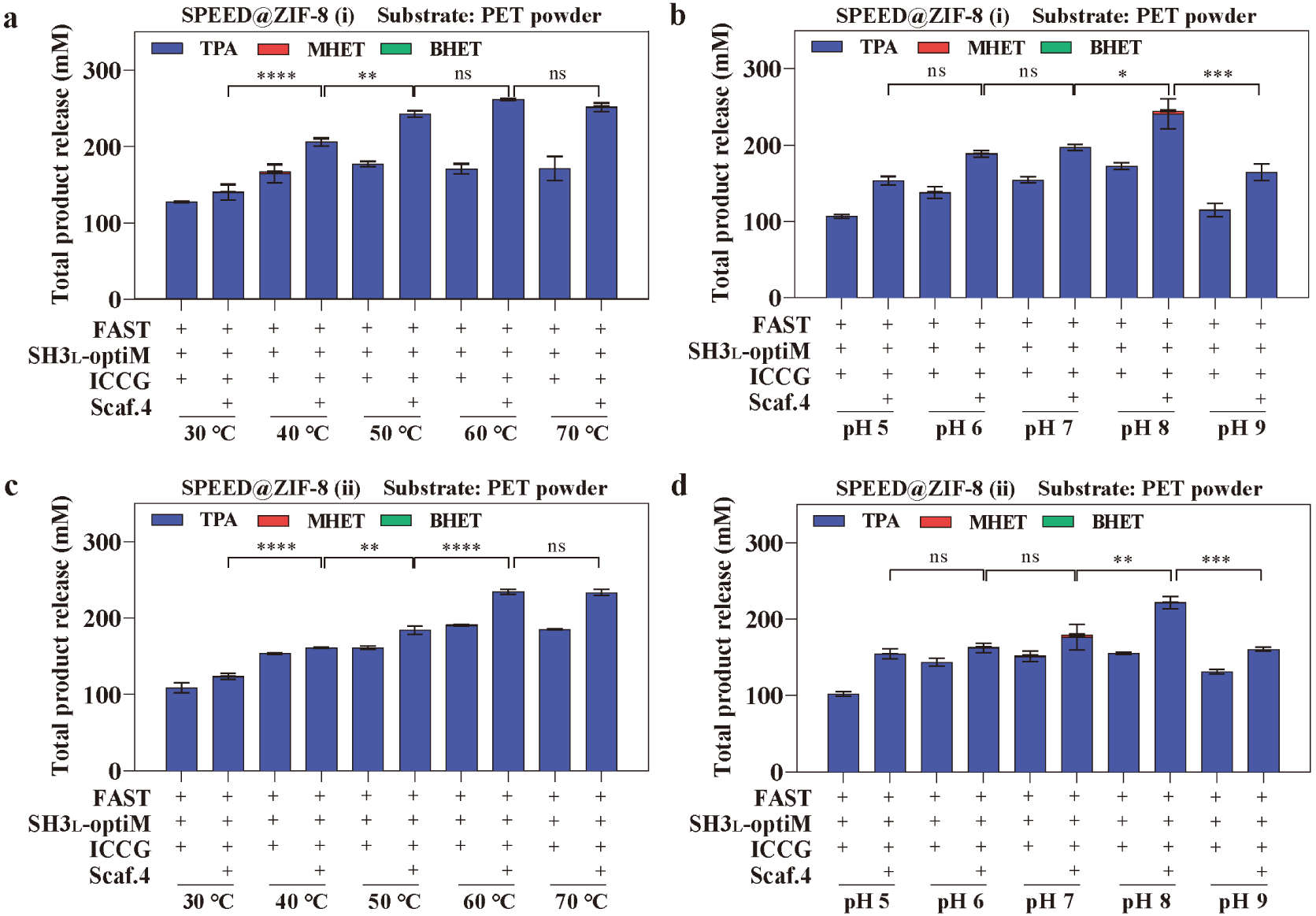
ZIF-8 immobilization broadens the operational window of optiComplex-D. Temperature (**a, c**) and pH (**b, d**) stability profiles for optiComplex-D immobilized on the surface of ZIF-8 (**a, b**) or via incomplete encapsulation (**c, d**). Compared to the free enzyme (Fig. 4), immobilization confers enhanced stability, particularly at higher temperatures. Reactions were performed with 100 mg PET powder, 0.25 nmol enzymes in a 0.5:1 enzyme:scaffold ratio for 96 h. The residual activity observed at low pH may be attributed to partial enzyme release from the ZIF-8 matrix under acidic conditions^64,65^. Data are presented as mean ± s.d. (n = 3). Statistical significance was determined by one-way ANOVA (ns: not significant, *p < 0.1, **p < 0.01, ***p < 0.001, ****p < 0.0001).

**Supplementary Fig. 14:**
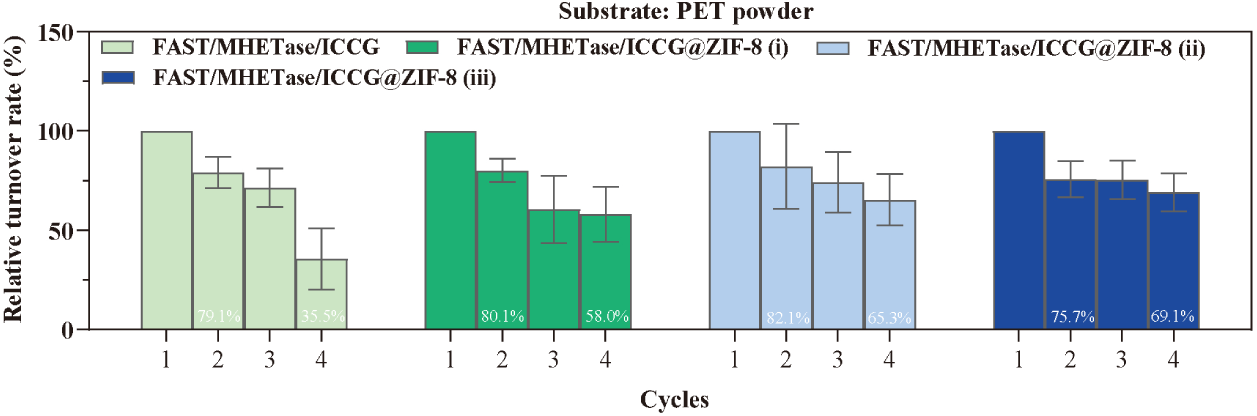
Reusability of ZIF-8 immobilized non-scaffolded enzymes. Relative activity of free and immobilized non-scaffolded enzyme mixtures (FAST/MHETase/ICCG) over four 4-day reaction cycles. Immobilization improves reusability compared to the free enzymes, but performance is inferior to the scaffolded complexes shown in Fig. 5d. Reactions were performed at 40 °C, pH 7.0 with 100 mg PET powder, 0.25 nmol individual enzymes. Data are presented as mean ± s.d. (n = 3).

**Supplementary Fig. 15:**
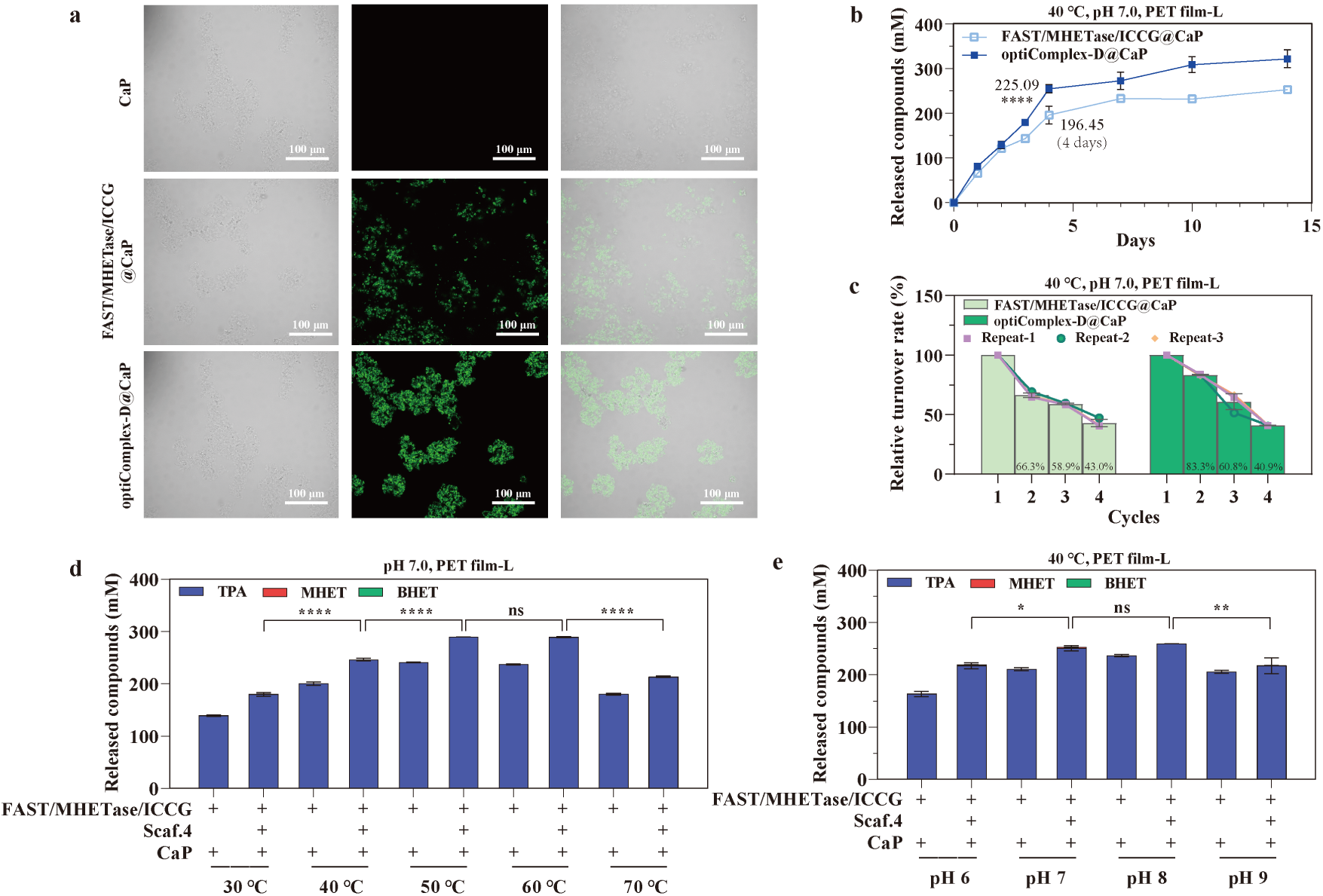
Performance of optiComplex-D immobilized on calcium phosphate (CaP). **a** Confocal microscopy images confirming successful immobilization of scaffolded (optiComplex-D) and non-scaffolded enzymes on CaP crystals. Alexa Fluor® 488 conjugated anti-His antibody was used to detect His-tagged proteins. The loading efficiency of FAST/MHETase/ICCG and optiComplex-D on CaP was 83.20% and 79.95%, respectively. **b** Long-term (14-day) time-course PET degradation analysis, showing an improvement over free enzyme complex. **c** Reusability over four 4-day cycles. FAST/MHETase/ICCG, a mixture of FAST-PETase, ICCG, and MHETase (0.25 nmol of each). **d**, **e** Temperature (**d**) and pH (**e**) stability profiles, demonstrating that CaP provides limited thermal and acid stability compared to ZIF-8 (**Supplementary Fig. 12**). Reactions were performed with 100 mg PET film-L using 0.5:1 enzyme:scaffold ratio. Data are presented as mean ± s.d. (n = 3). Statistical significance was determined by one-way ANOVA (ns: not significant, *p < 0.1, **p < 0.01, ***p < 0.001, ****p < 0.0001.)

**Supplementary Fig. 16:**
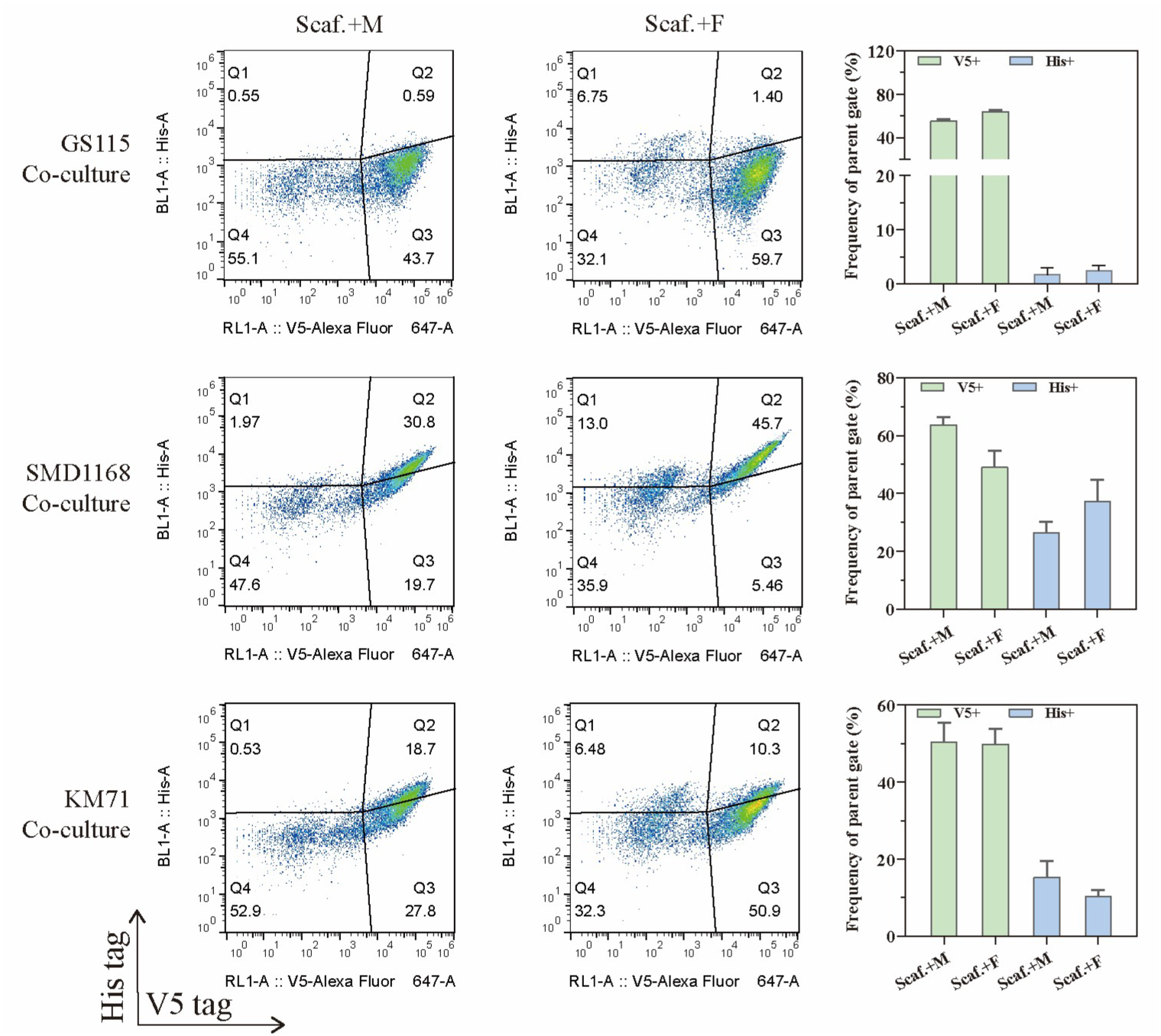
Screening of *P. pastoris* strains for scaffold expression and enzyme secretion. Flow cytometry analysis of co-cultures of V5-tagged scaffold-displaying and His-tagged enzyme-secreting yeast at 30℃ in BMMY medium. Three different *P. pastoris* host strains (GS115, SMD1168, KM71) were tested. Each co-culture consisted of a scaffold-displaying strain (Scaf.; His4::CipA-CBM-CipC-SED1) mixed at a 1:1 ratio with either an MHETase-secreting strain (M; His4::MHETase) or a FAST-secreting strain (F; His4::FAST). The SMD1168 strain showed the highest percentage of cells positive for both scaffold display and enzyme binding (co-localization), and was therefore selected for the experiments in Fig. 7.

### Supplementary Methods

#### Calculation of total enzyme molar amount

To facilitate a standardized comparison across different studies, the total molar amount of enzymes used in each reaction was calculated. Initial data, including either the mass or concentration of each enzyme, were extracted from the corresponding publications (**Supplementary File 1**). The total enzyme amount was determined by summing the molar amounts of the individual enzymes involved in each reported system. The molar amount for a single enzyme was calculated based on one of the following two scenarios:

1. For studies reporting enzyme mass, the molar amount was calculated as:

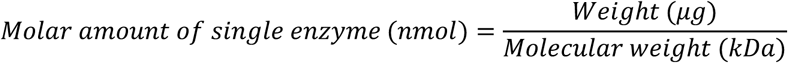 The molecular weight of each enzyme was determined using the ExPASy ProtParam tool.
2. For studies reporting enzyme concentration, the molar amount was calculated as:

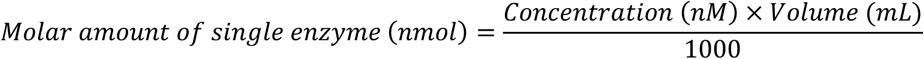

#### Calculation of total product molar amount

The total molar amount of degradation products (including TPA, EG, MHET, and BHET) was determined from product concentration or substrate mass loss data reported in the referenced literature (**Supplementary File 1**).

1. When product concentrations were provided, the total molar amount was calculated as:

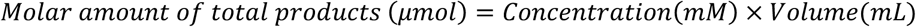
2. When substrate mass loss was provided, the total molar amount was calculated as:

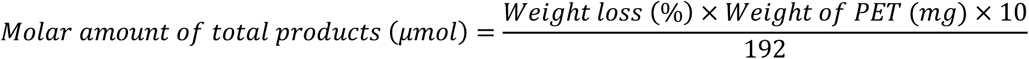

where 192 g/mol is the molecular weight of the polyethylene terephthalate (PET) repeating unit.

#### Calculation of degradation efficiency

To normalize catalytic performance, degradation efficiency was calculated as the total moles of product generated per mole of enzyme used.

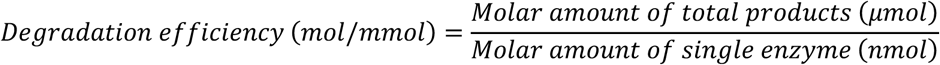

#### Calculation of monomer conversion fidelity

The efficiency of converting intermediates into the final recyclable monomers was assessed by calculating the percentage of monomer release. This metric reflects the fidelity of the multi-enzyme cascade.

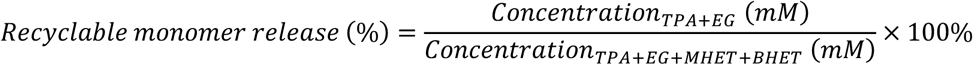

#### Calculation of PET conversion extent

The extent of PET substrate conversion was calculated based on the total molar amount of soluble degradation products (TPA, MHET, BHET) quantified by HPLC. The conversion percentage was determined using the following formula:

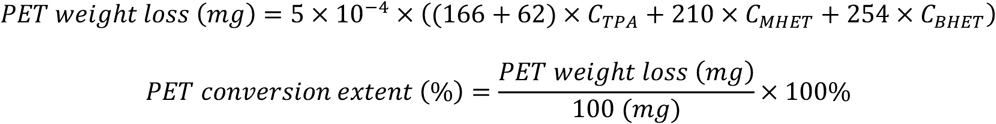

where C_TPA_, C_MHET_, C_BHET_ are the molar concentrations (mol/L) of TPA, MHET, and BHET, respectively. Note that EG has equal molar amount to TPA.

