## Supplementary File 1 for "Synergistic Multi-Enzyme Cascades Assembled on Modular Protein Scaffolds for Efficient PET Biorecycling"

**Supplemental File 1-1: Single-enzyme systems toward PET plastic biodegradation (associated with Supplementary Table 1)**

| Degradation system | Substrate | Brand of substrate | Crystallinity of substrate | Reaction condition | MW (kDa) | Substrates (mg PET) | Total Enzyme (μg) | Enzyme (nM) | Total enzyme (nmol) | Concentration of total products (mM) | System volume (mL) | Total products (umol) | Degradation efficiency (mol total products/mmol enzyme) | Data sourse | Reference |
|---|---|---|---|---|---|---|---|---|---|---|---|---|---|---|---|
| **IsPETase** | PET film | N/A | 2% | 30 °C, 18h, pH 7.0 | 29 | N/A | | 50 | 0.015 | 0.3 | 0.3 | 0.09 | 6.00 | Figure 2d | Yoshida et al., 2016 |
| **TfH** | PET film | | | 30 °C, 18h, pH 7.0 | 29 | N/A | | 50 | 0.015 | 0.00079 | 0.3 | 0.000237 | 0.02 | | |
| **LCC** | PET film | | | 30 °C, 18h, pH 7.0 | 29 | N/A | | 50 | 0.015 | 0.0063 | 0.3 | 0.00189 | 0.13 | | |
| **FsC** | PET film | | | 30 °C, 18h, pH 7.0 | 24 | N/A | | 50 | 0.015 | 0.00075 | 0.3 | 0.000225 | 0.02 | | |
| **LCC** | Post-consumer coloured-flake PET waste | N/A | 29% | 72 °C, 20 h, pH 8.0 | 29 | 16000 | | 8625 | 690 | 552.09 | 80 | 44167.2 | 64.01 | Figure 4a | Tournier et al., 2020 |
| **ICCG (LCC$^{F243I/D238C/S283C/Y127G}$)** | | | | 72 °C, 15 h, pH 8.0 | 29 | 16000 | | 8625 | 690 | 854.17 | 80 | 68333.6 | 99.03 | | |
| **WCCG (LCC$^{F243W/D238C/S283C/Y127G}$)** | | | | 72 °C, 20 h, pH 8.0 | 29 | 16000 | | 8625 | 690 | 885.42 | 80 | 70833.6 | 102.66 | | |
| **DuraPETase (IsPETase$^{S214H/I168R/W159H/S188Q/R280A/A180I/G165A/Q119Y/L117F/T140D}$)** | PET film | Artificially prepared | 30% | 50 °C, 72 h, pH 9.0 | 29 | 1.00 | 5.00 | 347.67 | 0.17 | 2.63 | 0.50 | 1.32 | 7.56 | Figure 3a, and Figure S8 | Cui et al., 2021 |
| | | | | 37 °C, 72 h, pH 9.0 | 29 | 1.00 | 5.00 | 347.67 | 0.17 | 1.76 | 0.50 | 0.88 | 5.06 | | |
| **LCC** | PET film | | | 37 °C, 72 h, pH 9.0 | 29 | 1.00 | 5.00 | 346.75 | 0.17 | 1.04 | 0.50 | 0.52 | 3.00 | | |
| **TS-PETase (IsPETase$^{S121E/D186H/R280A}$)** | PET film | Goodfellow, ES301445 | N/A (low-crystallinity, 250 μm in thickness, 18% approximately) | 30 °C, 6 days, pH 9.0 | 29 | N/A | | 400.00 | 0.40 | 0.05 | 1.00 | 0.05 | 0.13 | Figure S8 | Zhong-Johnson et al., 2021 |
| ***Thermobifida fusca* cutinase$^{H184S/Q92G/F209I/I213K}$** | PET bottle | N/A | N/A (high-crystallinity) | 60 °C, 96 h, pH 8.0 | 34 | 1000.00 | 50000.00 | N/A | 1461.18 | N/A | N/A | 3542.00 | 2.42 | Figure 10 | Chen et al., 2022 |
| **LCC$^{H184S/Q92G/F209I/I213K}$** | PET bottle | | | 60 °C, 96 h, pH 8.0 | 29 | 1000.00 | 50000.00 | N/A | 1742.22 | N/A | N/A | 1354.00 | 0.78 | | |
| **BhrPETase$^{H184S/Q92G/F209I/I213K}$** | PET bottle | | | 60 °C, 96 h, pH 8.0 | 29 | 1000.00 | 50000.00 | N/A | 1724.14 | N/A | N/A | 3802.00 | 2.21 | | |
| **HotPETase** | PET powder | Goodfellow | 30% | 60 °C, 5 h, pH 9.2 | 28 | 20.00 | 5.80 | 500.00 | 0.21 | 6.07 | 5.00 | 30.35 | 146.52 | Extended Data Figure 6 | Bell et al., 2022 |
| **DuraPETase-4M (DuraPETase$^{N233C/S282C/H214S/S245R}$)** | PET powder | Kexinda Technology Co. Ltd., China | N/A (high-crystallinity) | 60 °C, 96 h, pH 9.0 | 29 | 25.38 | 87.50 | 347.38 | 3.04 | 15.80 | 8.75 | 138.25 | 45.48 | Figure 4d | Liu et al., 2022 |
| **DuraPETase** | PET powder | | | 60 °C, 96 h, pH 9.0 | 29 | 25.38 | 87.50 | 347.67 | 3.04 | 5.70 | 8.75 | 49.88 | 16.39 | | |
| **FAST-PETase (*Is* PETase$^{S121E/D186H/R224Q/N233K/R280A}$)** | PET film | Goodfellow, US, 577-529-50 | N/A (high-crystallinity, 40-70% approximately) | 50 °C, 96 h, pH 8.0 | 31 | 11.40 | | 200.00 | 0.12 | 33.80 | 0.60 | 20.28 | 169.00 | Figure 1c, and Figure S2b | Lu et al., 2022 |
| **ICCM (LCC$^{F243I/D238C/S283C/N246M}$)** | PET film | | | 40 °C, 96 h, pH 8.0 | 29 | 11.40 | | 200.00 | 0.12 | 14.75 | 0.60 | 8.85 | 73.75 | | |
| | | | | 50 °C, 96 h, pH 8.0 | | 11.40 | | 200.00 | | 29.50 | 0.60 | 17.70 | 147.50 | | |
| **DSI- Tfcutinase$^{D204C/E253C}$** | PET particle | Goodfellow, Germany | 8% | 70 °C, 96 h, pH 8.0 | 33 | 200.00 | | 7000.00 | 14.00 | 300.00 | 2.00 | 600.00 | 42.86 | Figure 4 | Z. Liu et al., 2022 |
| ***Is* PETase-Spy (Cyclic monomer)** | PET powder | Goodfellow | N/A (Semi-crystalline) | 30 °C, 24 h, pH 7.5 | 43 | N/A | | 100.00 | 0.10 | 0.40 | 1.00 | 0.40 | 4.03 | Figure 7 | Hayes & Luk, 2023 |
| ***Is* PETase-Dimer (Cyclic dimer)** | PET powder | | | 30 °C, 24 h, pH 7.5 | 55 | N/A | | 50.00 | 0.05 | 0.46 | 1.00 | 0.46 | 9.16 | | |
| ***Is* PETase-Cat (Catenane)** | PET powder | | | 30 °C, 24 h, pH 7.5 | 50 | N/A | | 50.00 | 0.05 | 0.50 | 1.00 | 0.50 | 9.95 | | |
| ***Is* PETase** | PET powder | | | 30 °C, 24 h, pH 7.5 | 33 | N/A | | 100.00 | 0.10 | 0.40 | 1.00 | 0.40 | 3.95 | | |
| **TurboPETase** | PET film | Goodfellow, ES301445 | 7% (250 μm in thickness) | 65 °C, 12 h, pH 8.0 | 28 | 15.00 | 4.50 | 324.16 | 0.16 | 130.00 | 0.50 | 65.00 | 401.04 | Figure S7 | Cui et al., 2024 |
| **BhrPETase** | PET film | | | 65 °C, 12 h, pH 8.0 | 28 | 15.00 | 4.50 | 323.68 | 0.16 | 6.00 | 0.50 | 3.00 | 18.54 | | |
| **LCC$^{ICCG}$** | PET film | | | 65 °C, 12 h, pH 8.0 | 28 | 15.00 | 4.50 | 325.64 | 0.16 | 34.00 | 0.50 | 17.00 | 104.41 | | |
| **FAST-PETase** | PET film | | | 65 °C, 12 h, pH 8.0 | 31 | 15.00 | 4.50 | 290.32 | 0.15 | 2.00 | 0.50 | 1.00 | 6.89 | | |

**Green**: information from the corresponding publication.

**Black**: caculated numbers using the published information (Green).

**N/A**: information not available from the corresponding publication.

**Supplemental File 1-2: Multi-enzyme systems toward PET plastic biodegradation (associated with Supplementary Table 2)**

| Degradation system | Substrate | Brand of substrate | Crystallinity of substrate | Reaction condition | ratio of enzymes | MW (kDa) | Weight of substrate (mg) | Total Enzymes (μg) | Enzyme (nM) | Total enzymes (nmol) | Concentration of total products (mM) | System volume (mL) | Total products (umol) | Improved fold (comparing to single enzyme) | Degradation efficiency (mol total products/mmol enzyme) | Data sourse | Reference |
|---|---|---|---|---|---|---|---|---|---|---|---|---|---|---|---|---|---|
| **TfCa (immobilized) +LCC** | PET film | Goodfellow, Germany, 029-198-54 | N/A (low-crystallinity, 250 μm in thickness, 18% approximately) | 60 °C, 24 h, pH 9 | 3:1 | 55 of TfCa<br>29 of LCC | 6500.00 | 1600.00 | 541.96 | 35.55 | 10.42<br>6.71 (TPA) | 40.00 | 416.80<br>268.40 | 2.4 (to TfCut2 with TfCa) | 11.72<br>7.55 | Figure 3b. | Barth et al., 2016 |
| **HiC+CALB** | PET bottle | Crystal | 37% (0.1 mm in thickness) | 60 °C, 14 days, pH 7 | 9:1 | 32 of HiC<br>33 of CALB | 200.00 | 8000.00 | 49848.48 | 249.24 | 2.25 | 5.00 | 11.24 | 2.3 (to HiC) | 0.045 | Figure 3 | Carniel et al., 2017 |
| **HiC+CALB** | PET bottle | Crystal | 37% (0.1 mm in thickness) | 60 °C, 14 days, pH 7 | 1:1 | 32 of HiC<br>33 of CALB | 200.00 | 4000.00 | 6060.61 | 123.11 | 0.73 | 10.00 | 7.28 | 2.2 (to HiC) | 0.06 | Figure 3 | de Castro et al., 2017 |
| **PETase+MHETase** | PET film | Goodfellow, UK | 2%-3% | 30 °C, 96 h, pH 7.5 | 2:1 | 30 of PETase<br>62 of MHETase | 5.00 | 15.00 | 827.20 | 0.41 | 2.87 | 0.50 | 1.44 | 2.17 (to PETase) | 3.47 | Table S4 | Knott et al., 2020 |
| **cPETase** | PET film | Artificially prepared | 27% | 30 °C, 96 h pH 7.0 | | 30 | N/A | 300.00 | N/A | 10.00 | 0.01 | N/A | N/A | | N/A (0.643 M/mmolenzyme) | Figure 4b, and Figure 6a | Hwang et al., 2022 |
| **cCALB** | PET film | | | 30 °C, 96 h pH 7.0 | | 33 | N/A | 330.00 | N/A | 10.00 | 0.01 | N/A | N/A | | N/A (1.12 M/mmolenzyme) | | |
| **PC enzyme complex (cPETase+cCALB+scaffold)** | PET film | | | 30 °C, 96 h pH 7.0 | 1:1:1 | 63 of cPETase with CALB | N/A | 1260.00 | 500.00 | 20.00 | 0.09 | 40.00 | 3.55 | 13.8-fold (to cPETase) 7.9-fold (to | 0.18 | | |
| | Waste | | 34% | 30 °C, 96 h pH 7.0 | 1:1:1 | | N/A | 1260.00 | 500.00 | 20.00 | 0.04 | 40.00 | 1.60 | 3.50-fold (to CALB) | 0.08 | | |
| **LCC+TTCE** | PET film | Goodfellow, GF25214475 | 13% (0.25 mm in thickness) | 60 °C, 96 h, pH 8 | 1:1 | 30 of LCC<br>29 of TTCE | 200.00 | 58.80 | 4000.00 | 2.00 | 17.97 | 0.50 | 8.99 | 1.22 (to LCC) | 4.49 | Figure 5b | Mrigwani et al., 2022 |
| **IsPETasePM** | PET film | Goodfellow, ES30-FM-0001445 | N/A (low-crystallinity, 18% approximately) | 45 °C, 24 h, pH 7.5 | | 23 | 60.00 | | 50.00 | 0.05 | 0.40 | 1.00 | 0.40 | 4.2 (to IsPETasePM) | 8.00 | Figure 4b | von Haugwitz et al., 2022 |
| **TfCaWA (TfCa $^{I69W/V376A}$) +IsPETasePM (IsPETase $_{S121E/D186H/R280A/N233C/S282C}$)** | PET film | | | 45 °C, 24 h, pH 7.5 | 11:1 | 55 of TfCa<br>23 of IsPETasePM | 60.00 | 31.40 | 600.00 | 0.60 | 1.68 | 1.00 | 1.68 | | 2.80 | | |
| **SPEED (FAST+ICCG+MHETase+ scaffold)** | PET powder | Xing-xiang new materials, China | 40.82% (high-crystallinity) | | 1:1:1:2 | 40 of FAST<br>39 of ICCG<br>66 of MHETase | 100.00 | 36.47 | 1500.00 | 0.75 | 143.76 | 0.50 | 71.88 | 4.11 (to FAST-PETase)<br>3.43 (to ICCG) | 95.84 | Figure 4e (4 days) | This study |
| | PET film | Goodfellow, ES30-FM-0001445 | 17.99% (low-crystallinity) | 40 °C, 96 h, pH 7.0 | | | 100.00 | 36.47 | | | 148.82 (total products) | 0.50 | 74.41 | 3.47 (to FAST-PETase) | 99.21 | Figure 4f (4 days) | |
| | | | | | | | | | | | 146.75 (TPA) | 0.50 | 73.38 | 2.89 (to ICCG) | 97.83 | | |
| | PET film | Goodfellow, ES30-FM-000250 | 43.31% (high-crystallinity) | | | | 100.00 | 36.47 | | | 3.23 | 0.50 | 1.62 | 4.01 (to FAST-PETase)<br>2.92 (to ICCG) | 2.15 | Figure 4g (4 days) | |
| | PET film | Goodfellow, ES30-FM-0001445 | 17.99% (low-crystallinity) | 40 °C, 24 h, pH 7.0 | | | 100.00 | 36.47 | | | 82.87 | 0.50 | 41.44 | | 55.25 | Figure 4f (1 day) | |
| | | | | | | | | | | | 189.85 | 0.50 | 94.93 | | 126.57 | Figure 4f (14 days) | |
| | PET film | Goodfellow, ES30-FM-000250 | 43.31% (high-crystallinity) | 40 °C, 14 days, pH 7.0 | | | 100.00 | 36.47 | | | 9.05 | 0.50 | 4.52 | | 6.03 | Figure 4g (14 days) | |

**Green**: information from the corresponding publication.

**Black**: caculated numbers using the published information (Green).

**N/A**: information not available from the corresponding publication.
